# Sequence-Dependent Backbone Dynamics of Intrinsically Disordered Proteins

**DOI:** 10.1101/2022.02.11.480128

**Authors:** Souvik Dey, Matthew MacAinsh, Huan-Xiang Zhou

## Abstract

For intrinsically disordered proteins (IDPs), a pressing question is how sequence codes for function. Dynamics serves as a crucial link, reminiscent of the role of structure in sequence-function relations of structured proteins. To define general rules governing sequence-dependent backbone dynamics, we carried out long molecular dynamics simulations of eight IDPs. Blocks of residues exhibiting large amplitudes in slow dynamics are rigidified by local inter-residue interactions or secondary structures. A long region or an entire IDP can be slowed down by long-range contacts or secondary-structure packing. On the other hand, glycines promote fast dynamics and either demarcate rigid blocks or facilitate multiple modes of local and long-range inter-residue interactions. The sequence-dependent backbone dynamics endows IDPs with versatile response to binding partners, with some blocks recalcitrant while others readily adapting to intermolecular interactions.

## Introduction

The sequence-structure-function paradigm has guided protein biophysics for many decades. Intrinsically disordered proteins (IDPs) account for 30% to 50% of proteomes and perform myriad cellular functions including signaling and regulation ^1, 2^. Many IDPs, including amyloid-*β* peptides (e.g., A*β*40), tau, and *α*-synuclein, are also central players in human diseases such as Alzheimer’s and Parkinson’s ^3, 4^. The lack of well-defined structures poses the fundamental question of how IDP sequences code for functions.

Many studies have shown that, for IDPs, dynamics serves as a crucial link between sequence and function, replacing the role ascribed to structure for structured proteins. For example, fast dynamics allows IDPs to rapidly adapt to target proteins in forming stereospecific complexes ^5^. Likewise, uncoupled dynamics between neighboring blocks of an IDP enables fast dissociation from its target protein, leading to a desired short lifetime for the signaling complex ^6^. The level of dynamic coupling between neighboring blocks may also dictate the competition between IDPs in binding to the same target protein ^7, 8^.

Sequence-dependent backbone dynamics on the ps-ns timescales of many IDPs has been characterized by NMR relaxation ^9–17^. Whereas relaxation properties of structured proteins can be easily interpreted by well-separated ns global motions and ps internal motions ^18^, dynamics of IDPs occur on many scales that are intricately linked and therefore analyzing the resulting NMR relaxation properties is not trivial. One approach is to fit the time correlation function of each backbone N-H bond vector to a sum of exponentials ^11–13^, but then one is still at a loss regarding the molecular origins of any sequence dependence of the backbone dynamics. Another approach is to incorporate results from molecular dynamics (MD) simulations ^14, 16^. Force fields for MD simulations have become accurate for predicting conformational properties ^19–21^ but, up to the recent past, have not been reliable for predicting dynamic properties ^14, 16^. Therefore MD results for relaxation rates or time constants had to be rescaled in order to match with NMR relaxation data, which unfortunately introduces uncertainty to MD-derived interpretations of backbone dynamics. For *α*-synuclein, both residues with small amplitudes (the NAC region, i.e., residues 62-95) and large amplitudes (e.g., Tyr39) on the slow timescale play important roles in fibrillation, but the connection between backbone dynamics and fibrillation is unclear ^14^. In A*β*40, residues Leu16-Phe20 have large amplitudes on the slow time scale, and noting that these residues form direct contact with the protofibril surface ^22^, it was speculated that the higher rigidity of these residues reduces the entropic cost for monomer-protofibril binding ^16^. Similarly, a block of residues around Lys311 of tau nucleates its aggregation ^23^; in the K18 fragment (residues 244-372) these residues exhibit restricted fast motions ^10^.

Recent advances in the implementation of force fields and water models in MD simulations have led to accurate predictions of not only conformational but also dynamic properties of IDPs ^24, 25^. We tested multiple force fields in simulations of the disordered N-terminal region (NT) of ChiZ ^25^, a member of the cell division machinery of *Mycobacterium tuberculosis*. The AMBER ff14SB force field ^26^ in conjunction with the TIP4P-D water model ^27^ predicted well not only the small-angle X-ray scattering (SAXS) profile and chemical shifts but also NMR relaxation rates. Two blocks of residues in the N-terminal half (N-half) of the 64-residue ChiZ-NT exhibit large amplitudes on the slow timescale, due to salt bridges, hydrogen bonds, cation-pi interactions, as well as polyproline II (PPII) formation. It was proposed that the N-half, rigidified by intramolecular interactions, would be recalcitrant to interactions with binding partners, whereas the C-terminal half (C-half) would readily adapt to binding partners. This model is validated in a subsequent study on the association of ChiZ with acidic membranes, showing that the C-half of NT is dominant in forming membrane contacts ^28^. Our study on ChiZ backbone dynamics further hinted that the thermostat needed for regulating temperature in MD simulations can distort dynamics, and correcting for such distortions can further improve agreement with NMR relaxation data ^25^. We have now developed a method for removing thermostat distortions of protein dynamics ^29^.

To define general rules governing sequence-dependent backbone dynamics, here we performed long MD simulations of eight IDPs with varying sequence lengths and extents of secondary structure formations (Table 1 and Fig. 1). The MD results were validated by a variety of experimental data, including SAXS profiles, chemical shifts, paramagnetic relaxation enhancements (PREs), and NMR relaxation rates. The time correlation function of each backbone NH bond vector was fit to a sum of three exponentials, with time constants of 3-8 ns (*τ*_1_), 1-2 ns (*τ*_2_), and 0.1-0.2 ns (*τ*_3_). Blocks exhibiting large amplitudes in slow dynamics (*τ*_1_) are rigidified by side chain-side chain interactions or secondary structures, whereas glycines promote fast dynamics (*τ*_3_) and either demarcate rigid blocks or facilitate multiple modes of side chain-side chain interactions. The sequence-dependent backbone dynamics endows IDPs with versatile response to binding partners, with some blocks recalcitrant while others readily adapting to intermolecular interactions.

**Figure 1.**
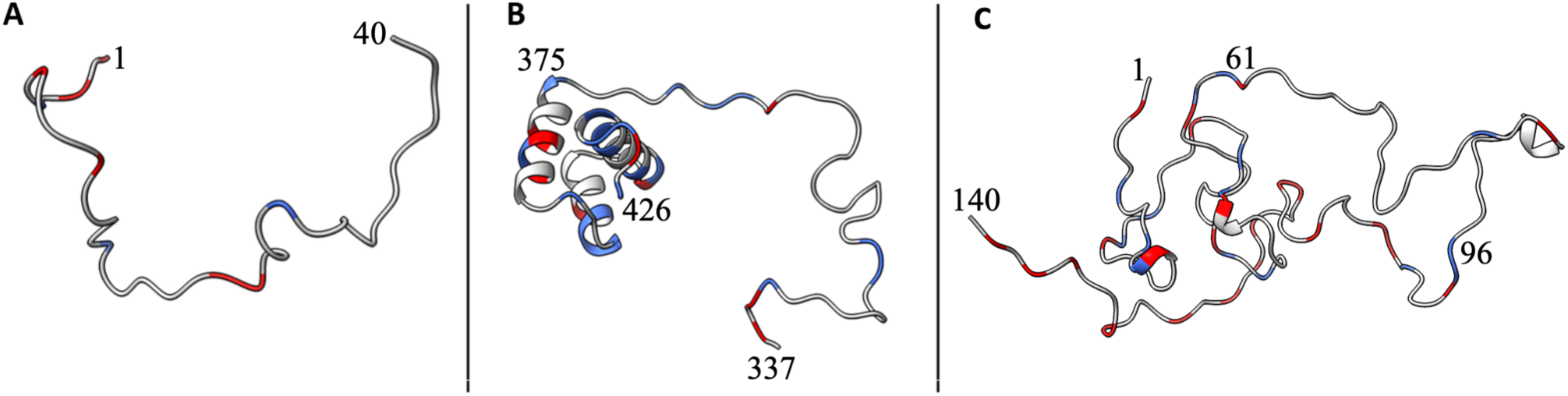
Snapshots from MD simulations of three IDPs. (A) Aβ-40. (B) HOX-DFD. (C) α-synuclein. Basic and acidic residues are colored blue and red, respectively. The N- terminal and C-terminal residue numbers are shown, as are the residue number marking the start of the helical region of HOX-DFD and the residue numbers marking the end of the N-terminal region and the start of the C-terminal region of α-synuclein.

**Table 1.**
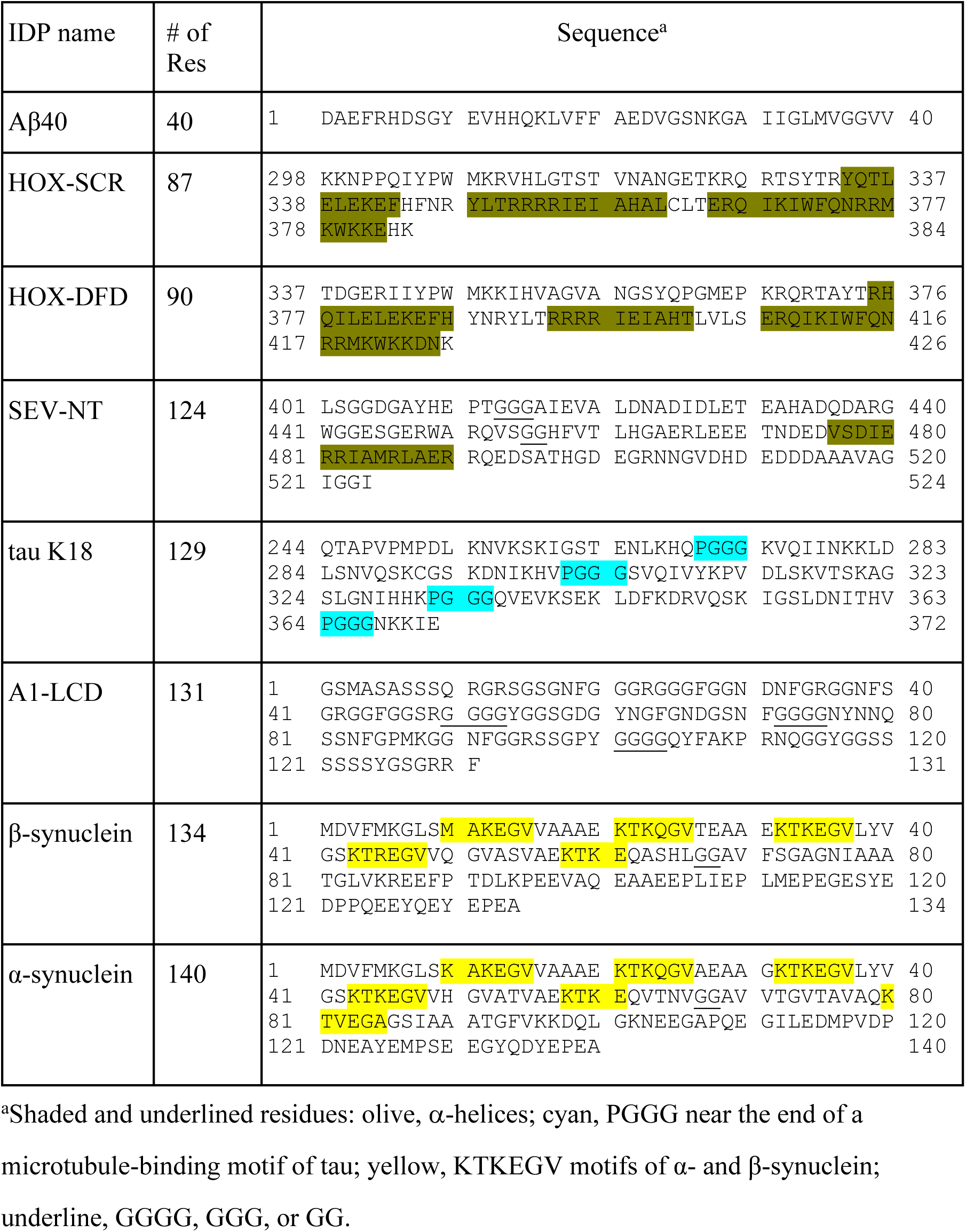
Sequences of eight IDPs.

## Results

The eight IDPs studied here range in sequence length from Aβ40 with 40 residues to α- synuclein with 140 residues (Table 1). Five of the proteins, Aβ40, tau K18, the low-complexity domain of heterogeneous nuclear ribonucleoprotein A1 (A1-LCD), α- and *β*- synuclein, are fully disordered ^10, 17, 30, 31^ (Fig. 1A,C), but two others, HOX transcription factors DFD and SCR, have three stable α-helices ^15^ (Fig. 1B), and one, the C-terminal domain of the nucleoprotein of Sendai virus (SEV-NT), has a well-populated α-helix ^11^. Following our previous study on ChiZ-NT ^25^, we used the AMBER ff14SB force field ^26^ and the TIP4P-D water model ^27^ to run four replicate simulations for each of the eight IDPs. Each replicate simulation was 3.2 μs, resulting in a total simulation time of 102.4 μs. The total numbers of atoms in the simulation systems ranged from 240,000 to 370,000 atoms (Table S1). Below we present validation of the MD results by a variety of experimental data (Table S1) and identify the origins of the sequence-dependent backbone dynamics.

### Experimental Validation of Conformational Ensembles From MD Simulations

SAXS profiles report on the overall size of a protein and have been determined for three of the IDPs, tau K18 ^32^, A1-LCD ^17^, and α-synuclein ^33^. In all the three cases, the SAXS profiles calculated on conformations sampled by our MD simulations show good agreement with the experimental counterparts (Fig. S1). To provide a direct measure of the sizes of the IDPs, we calculated the distributions of the radius of gyration (*R*_g_; Fig. S2). For three of the IDPs, Aβ40, tau K18, and α-synuclein, the mean *R*_g_ values agree with those predicted from a scaling relation,

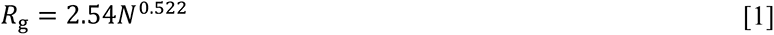

deduced from a set of IDPs (*N*: number of residues) ^34^. The mean *R*_g_ values of HOX-DFD, HOX-SCR, and SEV-NT are lower than predicted by Eq [1], due to the presence of α-helices. The mean *R*_g_ value of A1-LCD is also lower than predicted, due to extensive local and long-range interactions (see below).

Secondary chemical shifts indicate the formation of α-helices and *β*-strands, and have been measured for all the eight IDPs ^9–11, 15, 17, 30, 35^. In Figs. 2A-C and S3A-E, we compare the experimental data with the secondary chemical shifts calculated on conformations sampled by our MD simulations. Overall, there is reasonable agreement. Most of the secondary chemical shifts are close to 0, indicating complete disorder. The exceptions are three blocks in both of the HOX transcription factors and one block in SEV-NT, with large positive secondary chemical shifts indicating high propensities for α- helices. In later figures, we also report the helix and *β*-strand propensities of individual residues, which confirm the lack of well-populated secondary structures other than the α- helices in HOX-DFD, HOX-SCR, and SEV-NT.

**Figure 2.**
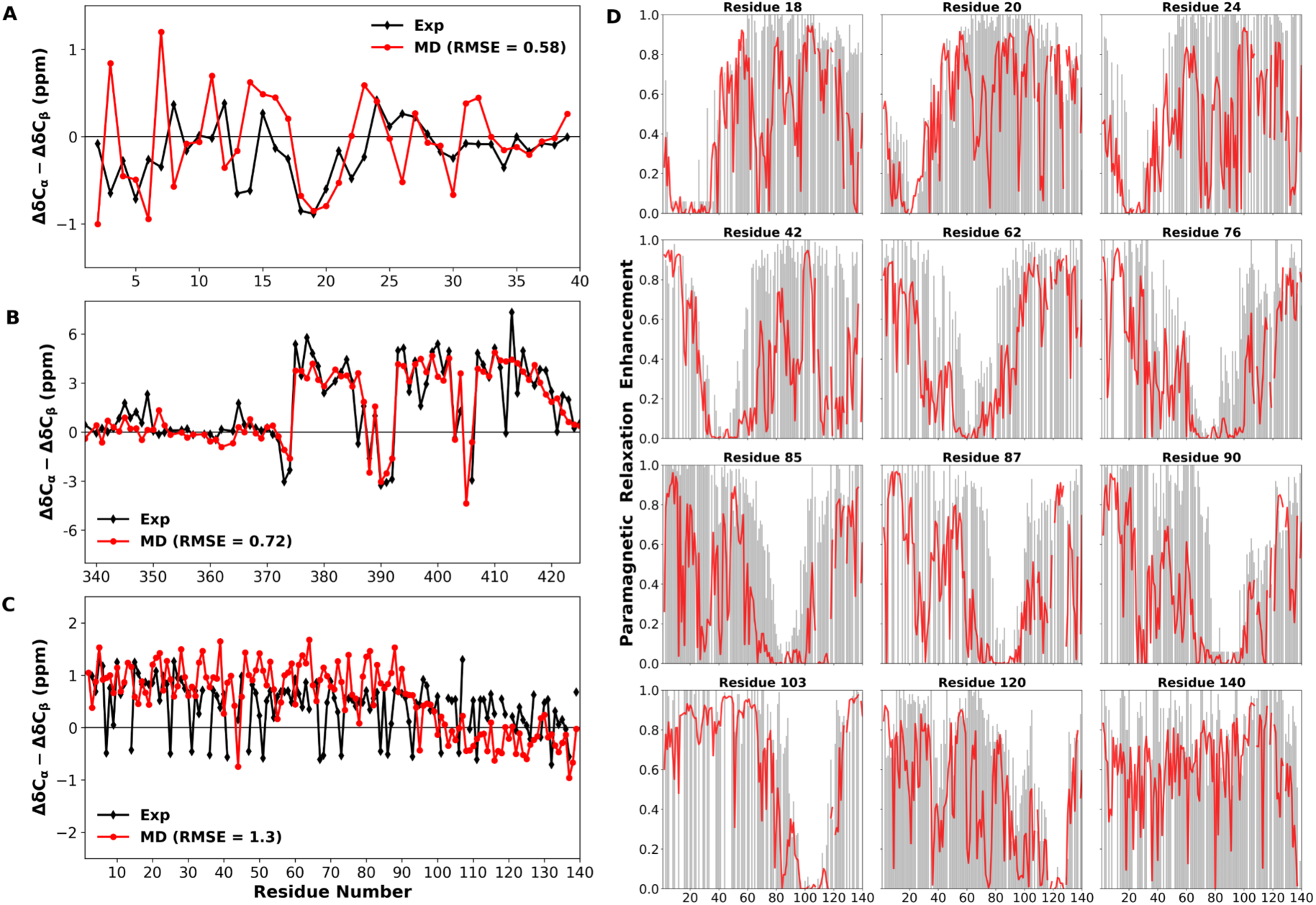
Experimental validation of MD conformational ensembles. (A-C) Comparison of calculated and experimental secondary chemical shifts for Aβ40, HOX-DFD, and α- synuclein. RMSE values are shown in the legends. (D) Comparison of calculated (red curves) and experimental (gray bars) PREs by spin labels at 12 residues in α-synuclein.

Inter-residue nuclear Overhauser effect (NOE) cross-peaks report on pairs of residues that can form close contact between hydrogen atoms; for Aβ40, NOEs between H_α_ at residue *i* and H_N_ at residue *i* + 2 “were seen within the Asp7-Glu11, Phe20-Ser26, and Gly29-Ile31 regions” ^30^ (Fig. S3F). We calculated the effective interproton distances, <r-^6^>^-1/6 36^, which implicated NOEs for His6, Asp7, His13, Asp23, and Asn27 (Fig. S3F). The calculated results, except for the addition of His13, are in agreement with the observed αN(*i*, *i* + 2) NOEs.

PREs can reveal long-range interactions, and have been measured for spin labels placed at 12 residues distributed throughout the sequence of α-synuclein ^31, 37–39^.

Consistent with the experimental data, a site in the N-terminal region (residues 1-61) can interact with the C-terminal region (residues 96-140) and vice versa; likewise a site in the central NAC region can interact with the C-terminal region and vice versa (Fig. 2D). For example, our MD results correctly predict that a spin label at residue 120 ^31^ has propensities for interacting with residues around position 40 and residues around position 90. The calculated PREs for *β*-synuclein do not agree as well with the experimental values ^31^ (Fig. S4A-C). The MD results correctly predict the absence of contacts between a spin label at position 20 with the central region, but incorrectly predict interaction propensities between a spin label at residue 114 with residues around position 10 and residues around position 75. For tau K18, consistent with experimental data ^40^, PREs calculated from the MD conformations indicate absence of long-range interactions (Fig. S4D). Next we document the validation of our MD simulations by NMR relaxation data.

### Correcting for Dynamic Distortion by the Langevin Thermostat

The backbone amide transverse (*R*_2_) and longitudinal (*R*_1_) relaxation rates and nuclear Overhauser effects (NOE) are determined by the time correlation function of the NH bond vector (see Computational Methods). We fit this correlation function to a sum of three exponentials, with time constants *τ*_1_, *τ*_2_, and *τ*_3_ (ordered from long to short) and amplitudes *A*_1_, *A*_2_, and *A*_3_ ^25^. Illustrative fits are shown in Fig. S5A-C. Note that we did not restrain the sum of the amplitudes to be 1; the missing amplitude represents ultrafast motion (faster than the 20-ps interval for saving snapshots from the MD simulations). The exponential components with the long (*τ*_1_), intermediate (*τ*_2_), and short (*τ*_3_) time constants contribute the most to *R*_2_, *R*_1_, and NOE, respectively.

We have recently shown systematic time dilation by the Langevin thermostat ^41^ used here and elsewhere for regulating temperature in MD simulations, and found a correction scheme ^29^. A raw time constant *τ*_raw_ obtained in an MD simulation is corrected by a scaling factor,

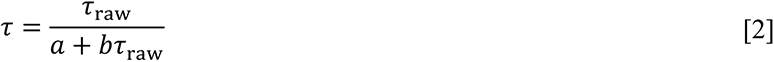

The constants *a* and *b* depend on the damping constant of the Langevin thermostat and are 1.526 and 0.086, respectively, for the damping constant of 3 ps^-1^ in the present study. The MD simulations in the present study were also run at constant pressure using the Berendsen barostat ^42^; however, we did not find any additional effect of the barostat on protein dynamics. Note that the scaling factor is proportional to *τ*_raw_, and thus has the strongest effect on *τ*_1_ and weakest effect on *τ*_3_. Figure S5D illustrates the correction for the three time constants of Gln15 in Aβ40 and of Ile397 in HOX-DFD. Figure S6 demonstrates that the correction for the thermostat effects dramatically reduces the root-mean-square-error (RMSE), from 3.1 s^-1^ to 0.5 s^-1^, of the *R*_2_ values of α-synuclein, while maintaining the sequence-dependent profile. In fact, the *R*_2_, *R*_1_, and NOE values calculated from the MD simulations match well with the experimental data for all the eight IDPs (Figs. 3 and S7A to S11A).

**Figure 3.**
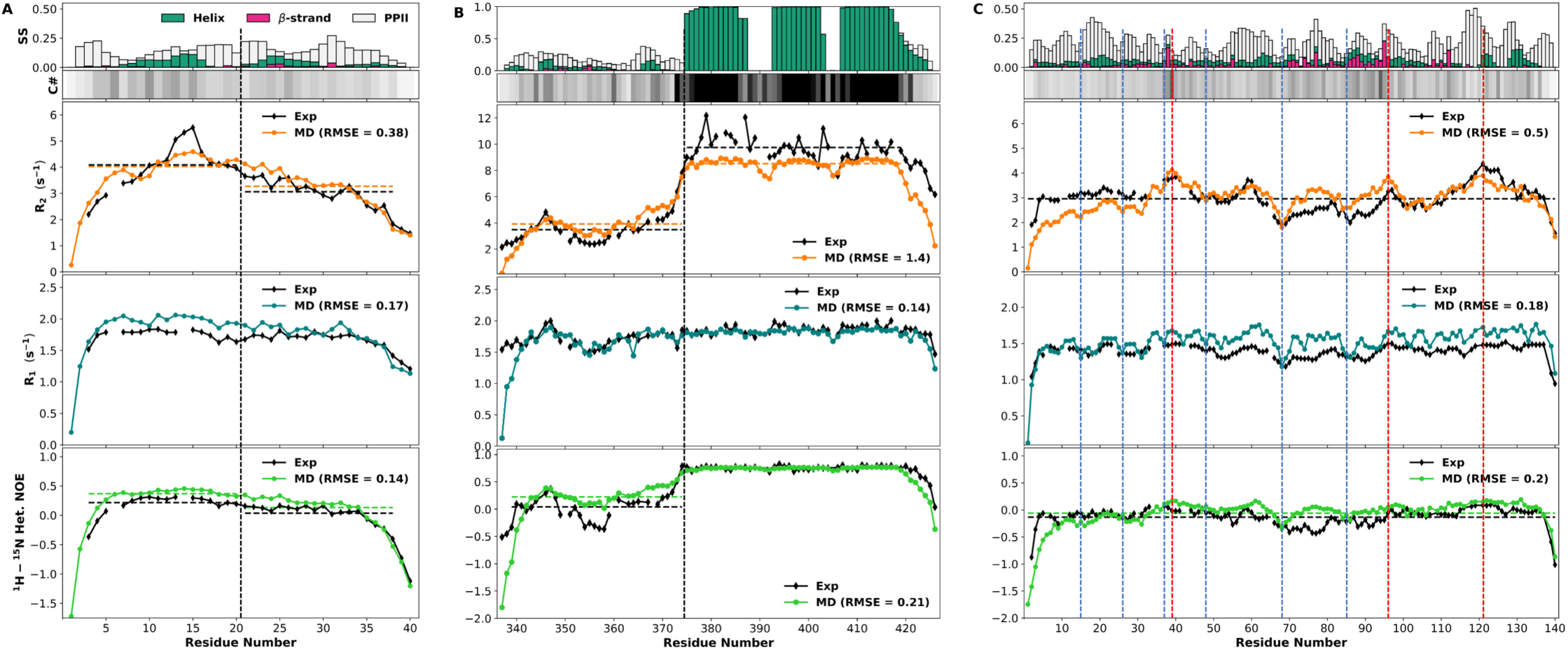
Backbone NMR relaxation properties. (A) Aβ40. (B) HOX-DFD. (C) α- synuclein. Calculated and experimental *R*_2_, *R*_1_, and NOE values are compared, with RMSE values shown in the legends. Horizontal dashed lines show average values over the N-half and C-half of Aβ40 or of HOX-DFD (demarcated by black vertical lines), or over the entire sequence of α-synuclein. For the latter protein, blue vertical lines are placed at residues 15, 26, 37, 48, 68, and 85 to indicate dips in *R*_1_ (and *R*_2_ in the case of Gly68); red vertical lines are placed at Tyr39, Lys96, and Asp121 to indicated elevated *R*_2_. Also shown at the top are secondary structure propensities and average contact numbers. The latter are displayed as bars on a gray scale, with white at 0 and black at 5 or above.

### Backbone Dynamics of Aβ40

Figure 3A displays the comparison between the calculated and experimental ^16^ *R*_2_, *R*_1_, and NOE values for Aβ40 residues. We make the following observations on the sequence dependences of the calculated results. First, the profile of each relaxation property along the sequence has a bell shape, with the terminal residues having much lower values than internal residues. This observation holds for all the IDPs, and the corresponding elevated fast dynamics of the terminal residues on the fast timescale can be attributed to the fact that these residues are linked to the polypeptide chain only on one side. Second, the N-half of the sequence has higher *R*_2_ and NOE values than the C-half; the mean *R*_2_ values are ∼4 s^-1^ and 3 s^-1^ for the N- and C-halves, respectively. This asymmetry, corresponding to more significant slow dynamics in the N-half, is similar to what was observed for ChiZ-NT ^25^. Third, the highest *R*_2_ values are found for residues His13-His14-Gln15. The experimental *R*_2_ values are even higher, likely due to exchange between unprotonated and protonated histidines ^16^ that is not modeled by our MD simulations. Fourth, a local *R*_2_ peak is observed in His6-Gln7 of the N-half, while the first block of eight residues in the C-half (Ala21-Lys28) has higher *R*_2_ values than the next block of eight residues.

To uncover the origins of the sequence-dependent backbone dynamics (of the non-terminal residues), we calculated the secondary structure propensities and contact numbers of individual residues (displayed above the relaxation properties in Fig. 3A). The latter was defined as the number of residues, other than the immediate neighbors, that form contacts with a given residue; contacts were defined as between heavy atoms within a cutoff distance of 3.5 Å. In line with the secondary chemical shifts (Fig. 2A), helices and *β*-strands are only sampled infrequently (< ∼ 10% in all residues). PPIIs are also only sampled at low levels. The lowly populated secondary structures in Aβ40 do not explain the sequence-dependent backbone dynamics. Instead, the contact numbers provide the explanations. It is immediately clear that the N-half residues have higher contact numbers than the C-half residues. Moreover, His6, His13, and Gln15 that are in the local or global *R*_2_ peaks are among the residues with the highest contact numbers.

To further characterize residue-residue contacts, we calculated the contact map, which displays the frequencies of contact formation between any pair of side chain heavy atoms ^25^. The contact frequencies above a threshold of 0.005 are displayed in Fig. 4A.

**Figure 4.**
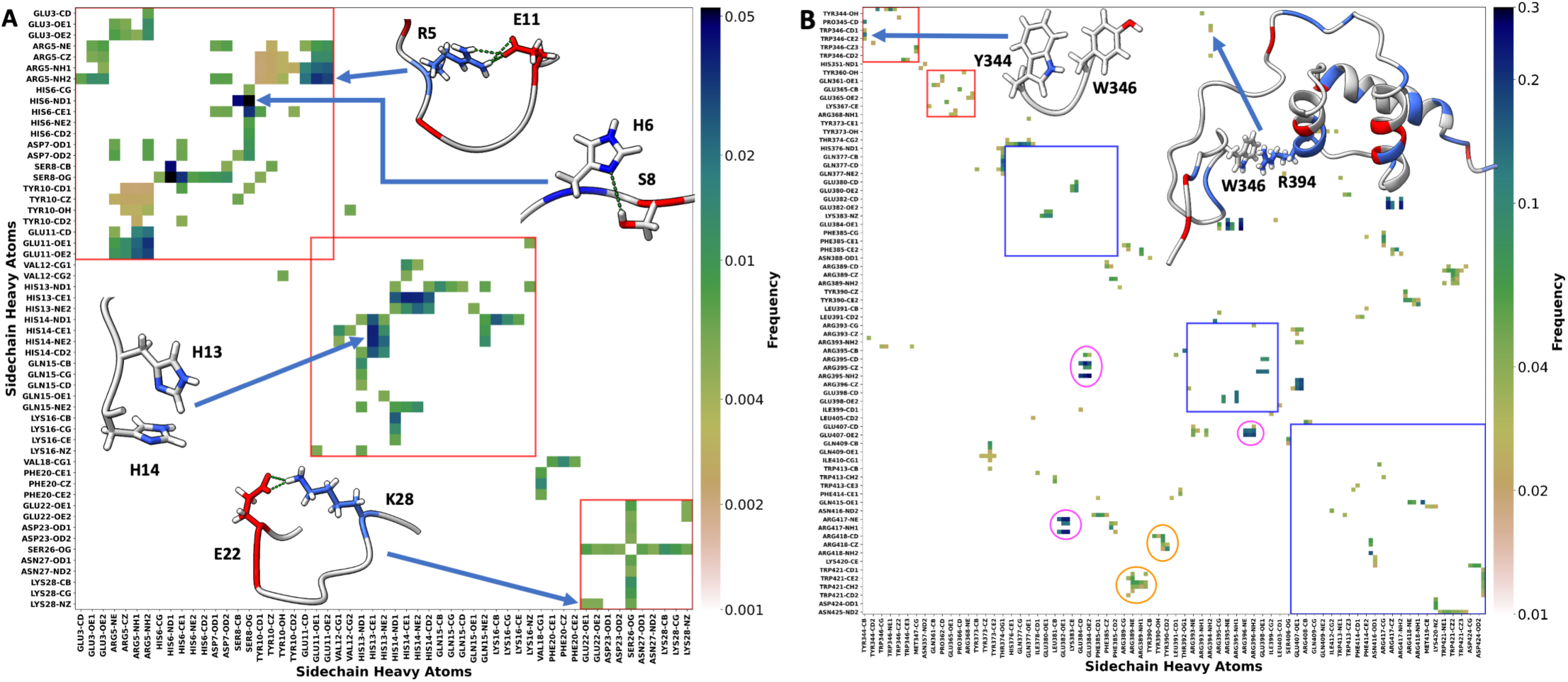
Side chain-side chain contact maps. (A) Aβ40. Contacts with frequencies higher than 0.005 are shown, but for Arg5-Tyr10 contacts, the threshold is lowered to 0.002. (B) HOX-DFD. Contacts with frequencies higher than 0.02 are shown. Blocks of residues with tendencies for extensive contacts are highlighted by red boxes. For HOX- DFD, α-helices are identified by blue boxes, and prominent interhelical and linker-helix contacts are indicated by magenta and orange ovals. Insets: snapshots illustrating selected contacts.

Two blocks of residues, Glu3- Glu11 and Glu11-Lys16, show tendencies to form extensive contacts, leading to rigidification. That these two blocks overlap and are both in the N-half underlies the asymmetry in backbone dynamics between the N- and C-halves. In the Glu3-Glu11 block, the most frequent contacts include salt bridges of Arg5 with Glu3, Asp7, and Glu11 (Fig. 4A inset); hydrogen bonds of Ser8 with His6 (Fig. 4A inset) and Asp7; and cation-π interaction between Arg5 and Tyr10. In the Glu11-Lys16 block, the most frequent contacts include a salt bridge of Glu11 with Lys16; π-π stacking between His13 and His14 (Fig. 4A inset); a hydrogen bond between His13 and Lys16; and amino-π interactions of Gln15 with His13 and His14. These side chain-side chain interactions explain the local *R*_2_ peak in residues 6-7 and the global *R*_2_ peak in residues 13-15. In the C-half, the Glu22-Lys28 block has a tendency to form a less extensive network of contacts, including a salt bridge between Glu22 and Lys28 (Fig. 4A inset) and hydrogen bonds of Ser26 with Glu22, Asp23, Asn27, and Lys28. These interactions explain why the first eight residues of the C-half have higher *R*_2_ values than the rest of the C-half. Note that the two serine residues, Ser8 and Ser26, can both hydrogen bond with multiple neighboring residues. These hydrogen bonds may be the reason for the αN(*i*, *i* + 2) NOEs described above.

All the foregoing interactions involve polar (including charged) residues. Val18 and Phe20 are the only two nonpolar residues that form contacts with elevated frequencies. The N-and C-halves contain 12 and 5 polar residues, respectively, and thus the former sequence is much more polar than the latter sequence. Therefore the origin of the asymmetry in backbone dynamics can be traced to the disparity in number of polar residues between the two halves.

### Backbone Dynamics of HOX-DFD

In Fig. 3B we compare the calculated and experimental ^15^ NMR relaxation data for HOX-DFD. In line with the experimental data, the most noticeable feature of the calculated relaxation data along the sequence is the significantly higher *R*_2_ and NOE of the helical C-half (starting at residue Arg375) relative to the disordered N-half. The mean *R*_2_ values are ∼4 s^-1^ and 9 s^-1^ for the disordered and helical regions, respectively. In contrast, *R*_1_ is relatively uniform, except for a dip around residue Gly354. In the helical region, *R*_2_ shows dips in the two inter-helical linkers. In the disordered region, *R*_2_, *R*_1_, and NOE all show a local peak in residues Trp346-Met347, which is a part of the conserved YPWM motif. After the dip around Gly354, *R*_2_ rises steadily from Tyr360 to Arg368, and then rapidly until reaching the helical region.

The high *R*_2_ and NOE of the C-half can be easily explained by the three α-helices, which remain intact in the MD simulations (Fig. 3B, top row). On the other hand, the N-half has low secondary structure content and forms few contacts. The contact numbers do show higher values in the YPWM motif and in the 14 residues (starting at Tyr360) preceding the helical region, and thus track well the sequence dependence of *R*_2_ in the N-half. The contact map, with a frequency threshold of 0.02, in Fig. 4B provides more details. In the C-half, 112 side-chain heavy atoms, or 41%, form contacts with other side-chain heavy atoms with frequencies above the threshold. The counterparts in the N-half are only 33 and 22%. The side chain-side chain contacts in the C-half show a well-packed 3-helix bundle, including salt bridges between the helices (Glu384-Arg395, Glu382- Arg417, and Arg396-Glu407) and cation-π interactions between the inter-helical linkers and the helices (Arg389-Trp421 and Tyr390-Arg418). In the N-half, most of the atoms with elevated contact frequencies belong to two blocks: the YPWM motif and Tyr360 to Arg368. The contacts in the YPWM motif are anchored by the Tyr344-Trp346 π-π stacking (Fig. 4B inset). In addition, this motif can occasionally form long-range contacts with the second α-helix in the C-terminal region, including a hydrogen bond between Tyr344 and Arg393 and a cation-π interaction between Trp346 and Arg394 (Fig. 4B inset). These interactions explain the local *R*_2_ peak in residues Trp346-Met347. The interactions within the Tyr360 to Arg368 block (e.g., a Glu365-Arg368 salt bridge) and between Tyr373 with the third α-helix and between Thr374 and the first α-helix explain the rise in *R*_2_ to the level of the C-terminal helical region.

### Backbone Dynamics of α-Synuclein

The calculated and experimental ^14^ *R*_2_, *R*_1_, and NOE values for α-synuclein are shown in Fig. 3C. In both sets of data, *R*_1_ and NOE are rather uniform along the sequence, but *R*_2_, with a mean value of ∼ 3 s^-1^, shows local peaks at Tyr39, Lys96, and Asp121. α-Synuclein contains six imperfect repeats with a consensus sequence KTKEGV (Table 1). In five of these repeats, the fifth position is a glycine. *R*_1_ dips at the sixth position in each of these five repeats (vertical lines in Fig. 3C). Two adjacent Gly residues occupy positions 67 and 68. Each of the three relaxation properties dips prominently at Gly68.

The secondary structure propensities and contact numbers provide some explanations for the *R*_2_ local peaks. Both Leu38 and Val95 have relatively high propensities (∼20%) for *β*-strands, while residues near Asp121 have PPII propensities close to 50%. Four proline residues, at positions 108, 117, 120, and 128, surround Asp121. Moreover, Tyr39 and those near Lys96 have the highest contact numbers. The contact map, with a frequency threshold of 0.005, provides more details (Fig. S12). Tyr39 is upstream of the fourth KTKE motif and can hydrogen bond with Ser42 (Fig. S12 inset); the latter can also form additional hydrogen bonds with Thr44 and Lys45 from the fourth KTKE motif.

These interactions explain the elevated *R*_2_ of Tyr39. Lys96 starts a block of residues with elevated frequencies for contact formation, including a salt bridge between Lys97 and Asp98 and hydrogen bonding of Lys96 with Gln99 (Fig. S12 inset) and of Asn103 with Gln 99 and Glu105. Furthermore, Lys96, along with Gln99, can also form long-range contacts with Asp121 (Fig. S12 inset). These interactions explain the elevated *R*_2_ of Lys96. In addition to the just-noted long-range contacts, the elevated *R*_2_ of Asp121 is also contributed by its location inside a block of residues, Pro108-Pro128, that contain four Pro residues and thus have high propensities for forming PPII.

### Backbone Dynamics of the Other Five IDPs

HOX-SCR is a very close homologue of HOX-DFD and their relaxation properties show very similar sequence dependences (Figs. 3B and S7A). Likewise β- and α-synuclein have significant sequence similarity, and their relaxation properties are all pretty uniform along the sequence, except for minor peaks in *R*_2_ (Figs. 3C and S8A). Compared to the HOX proteins and the synucleins, SEV- NT is intermediate both in the level of secondary structure formation and in the variation of relaxation properties over the sequence (Fig. S9A). Whereas the three α-helices of the HOX proteins are intact, the α-helix (Val476 to Gln492) in SEV-NT is only transient: the helical propensities peak at 90% over three residues but taper off on both sides. *R*_2_ exhibits a significant elevation in the middle of this transient helix, though the MD results do not rise as much as the experimental values ^11^. NOE also shows a modest rise whereas *R*_1_ shows a depression over the helical region. NOE and *R*_1_ are otherwise very uniform, but *R*_2_ shows a local peak at Ala450 and dips at Gly415 and Gly456. That glycines promote fast dynamics is now a familiar occurrence. The elevated *R*_2_ of Ala450 can be explained by its proximity to a block of residues, Gln436-Lys448, that tends to form extensive interactions, including Arg439-Asp437, Arg439-Glu444, and Arg448-Glu444 salt bridges, Gln436-Arg439 and Trp441-Glu444 hydrogen bonds, and Arg448-Trp441 cation-π interaction. Arg439 can also form salt bridges with upstream Asp427, Glu429, and Glu431, as well as Asp494 that is downstream of the transient helix. Glu429 can also form a salt bridge with Arg482 that is within the helical region. These long-range interactions help keep SEV-NT compact (Fig. S2D).

Tau K18 and Al-LCD happen to have similar sequence lengths (129 and 131 residues, respectively). While Al-LCD has propensities (∼20%) for β-strands in some residues, both of these IDPs lack stable secondary structures (Figs. S10A and S11A, top). However, as already noted above, whereas the overall size of tau K18 is typical of IDPs, Al-LCD is much more compact (Figs. S2E,F). Correspondingly, slow dynamics is largely absent in tau K18 but prominent in Al-LCD. In tau K18, *R*_2_ is below 5 s^-1^ for all but a few residues (e.g., Lys311 and Val313) (Figs. S10A). In contrast, in A1-LCD, *R*_2_ is above 5 s^-1^ for essentially all non-terminal residues, and reaches 9 s^-1^ for Tyr61-Phe64 and Asn79- Gln80 (Figs. S11A). The contact maps explain the difference. In tau K18, contacts with frequencies above 0.01 are sparse and all local (Fig. S13A). The sequence of this IDP can be divided into four imperfect repeats ending with PGGG (Table 1 and Fig. S13A). All the three relaxation properties show prominent dips at the end of each repeat (Figs. S10A). That observation, along with the lack of any significant contact between the repeats, suggests that the four repeats behave dynamically as independent units. In the third repeat, the relatively higher *R*_2_ values of Lys311 and Val313 can be accounted for by local contacts including a salt bridge between Lys311 and Asp314 and a hydrogen bond between Asp314 and Ser316 (Fig. S13A), and additionally by some rigidification provided by Pro312.

The sequence of Al-LCD contains three GGGG motifs, with the third Gly at positions 52, 74, and 103 (Table 1). *R*_2_ dips at these positions; the dips are particularly prominent for the last two GGGG motifs, where dips in NOE also occur (Figs. S11A). The contact map of Al-LCD shows three blocks of residues, Phe19-Arg49, Ser57-Phe71, and Asn76-Phe84, with elevated frequencies for extensive contacts (Fig. S13B). It is clear that the GGGG motifs demarcate these blocks. Within the second block, hydrogen bonds can be formed by Ser57 with Asp59, Asp59 with Asn62, Asn62 with Asn66, Asn66 with Asn70, and Asp67 with Ser69, and an amino-π interaction can be formed between Asn70 and Phe71. This block contains five Gly residues, and it appears that the flexibility of these Gly residues facilitates neighboring residues in forming multiple modes of interactions, e.g., Asp59 hydrogen bonding with either Ser57 or Asn62. In addition, residues in this block can also form long-range interactions, including hydrogen bonding of Tyr54 with Ser16, Asn62 with Gln113 and Asn79, and Asn66 with Asn91, cation-π interactions of Tyr61 (and Tyr54) with Arg95 (Fig. S13B inset) and of Phe64 with Arg95, and an amino-π interaction of Tyr61 with Asn91. Again, intervening Gly residues, e.g., between Asn91 and Arg95, appear to facilitate the multiple modes of interactions. These extensive networks of interactions explain the elevated *R*_2_ of Tyr61- Phe64. In the third block, hydrogen bonds can form between Asn76 and Asn78 or Gln80, between Ser81 and Asn83, and an amino-π interaction can form between Asn83 and Phe84, along with the already noted inter-block hydrogen bond between Asn62 and Asn79. These interactions explain the elevated *R*_2_ of Asn79-Gln80.

### Time Constants and Amplitudes of Three-Exponential Fits

Our NMR relaxation properties were calculated from the three-exponential fits of backbone NH time correlation functions. The time constants and amplitudes of these fits provide additional insight into the backbone dynamics of the IDPs. We display the distributions of the three time constants for each IDP in Fig. S14, showing good separation. The ranges of *τ*_1_, *τ*_2_, and *τ*_3_ are 3-8 ns, 1-2 ns, and 0.1-0.2 ns, respectively. The means and standard deviations of each time constant and the corresponding amplitude, calculated among all the residues of each IDP, are collected in Table S2; the values for individual residues are shown in Figs. 5 and S7B,C to S11B,C. The mean *τ*_1_ values separate the 8 IDPs into two groups.

**Figure 5.**
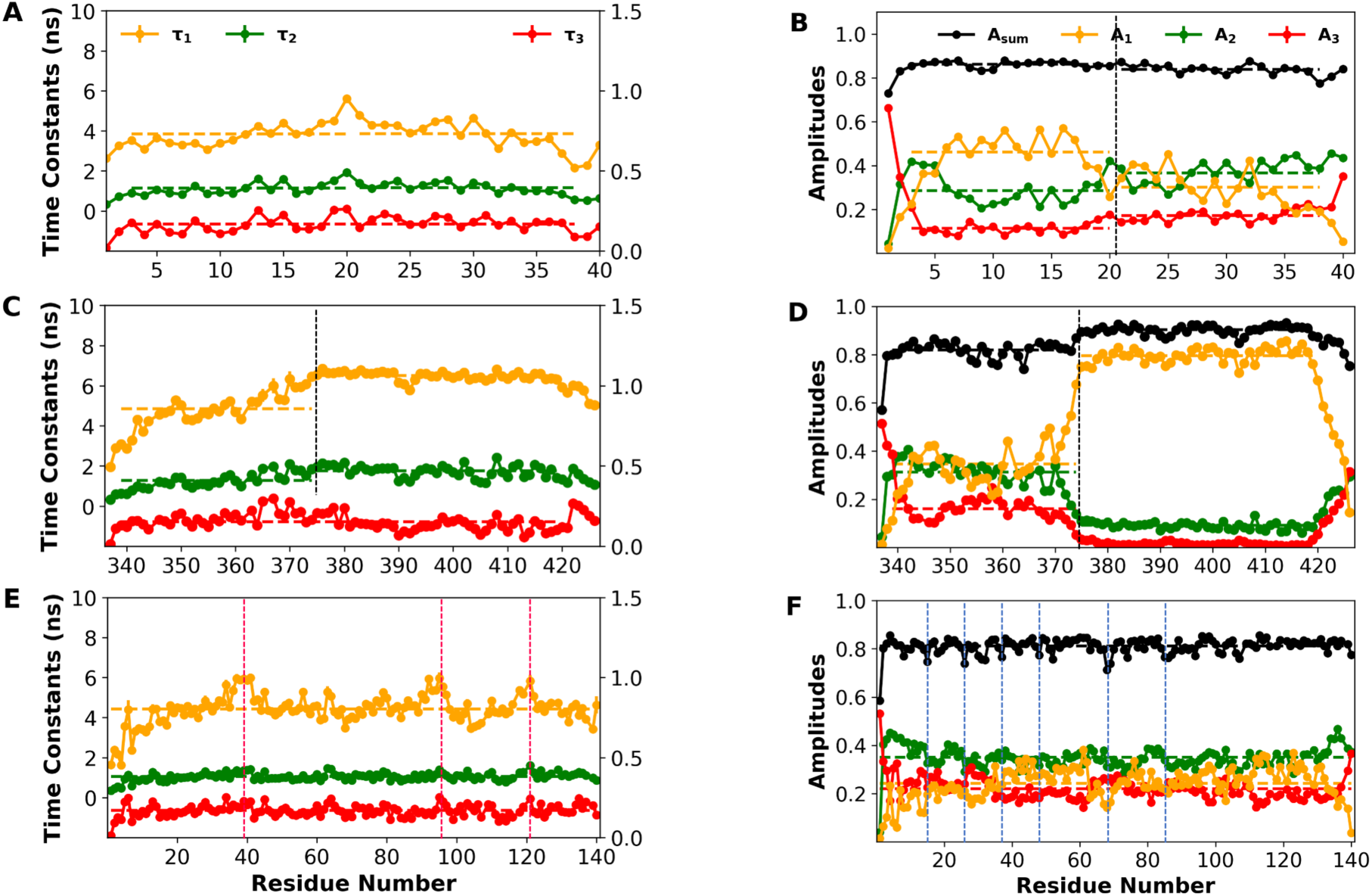
Time constants and amplitudes from fitting NH time correlation functions to a sum of three exponentials. (A-B) Aβ40. (C-D) HOX-DFD. (E-F) α-synuclein. In panels (A), (C), and (E), the left ordinate displays the scale for *τ*_1_ and *τ*_2_ whereas the right ordinate displays the scale for *τ*_3_. In panels (B), (D), and (F), the sum of the amplitudes, *A*_sum_, was not constrained to 1 and is shown in black; the missing amplitude is due to ultrafast motions. Horizontal dashed lines show average values over the N-half and C-half (demarcated by black vertical lines) or over the entire sequence. For α-synuclein, vertical lines are placed in panel (E) at Tyr39, Lys96, and Asp121 to indicate elevated *τ*_1_, and in panel (F) at residues 15, 26, 37, 48, 68, and 85 to indicate dips in *A*_sum_.

The first group, with mean *τ*_1_ close to or even above 6 ns, consists of HOX-DFD, HOX- SCR, and SEV-NT, all of which have stable or well-populated α-helices, and Al-LCD, which form numerous side chain-side chain contacts. The other four IDPs, with mean *τ*_1_ around 4 ns, are fully disordered and form much fewer side chain-side chain contacts than A1-LCD (β-synuclein has a mean *τ*_1_ at 5.2 ns but that is probably overestimated). The mean *τ*_2_ values of the first group are close to 1.6 ns and somewhat higher than those the second group, which are close to 1 ns. However, there is no distinction in mean *τ*_3_ between the two groups of IDPs. The first group has approximately double the mean *A*_1_ (at ∼0.6) but half the mean *A*_2_ (at ∼0.2) and mean *A*_3_ ( at 0.1) of the second group.

Correspondingly, *A*_1_ is the dominant amplitude in the first group but *A*_1_ and *A*_2_ are roughly equal in the second group. It is clear that the formation of secondary structures and side chain-side chain interactions promotes slow dynamics and suppresses fast dynamics.

Next we examine the sequence dependences of the time constants and amplitudes of the IDPs. Figures 5C and S7B show that, for HOX-DFD and HOX-SCR, the helical C-half has moderately higher *τ*_1_, slightly higher *τ*_2_, and similar *τ*_3_ when compared with the disordered N-half. More interestingly, as shown in Fig. 5D and S7C, going from the N-half to the C-half, there is a significant increase in *A*_1_ along with significant decreases in *A*_2_ and *A*_3_. Correspondingly, *A*_1_ and *A*_2_ are approximately equal in the N-half but *A*_1_ dominates in the C-half. The contrasts in time constants and amplitudes between the C- and N-halves of the two HOX proteins mirror precisely the contrasts between the IDPs as two separate groups, one rich in structures or interactions and the other lacking such features. The same contrasts apply to SEV-NT when the helical region is compared to the non-helical region, except that the dominance of *A*_1_ extends to most of the non-helical region as well (Fig. S9B,C), due to the long-range contacts detailed above. Likewise, due to the extensive local and long-range contacts, *A*_1_ dominates nearly the entire sequence of A1-LCD (Fig. S11B,C). The opposite scenario is represented by α-synuclein and tau K18, which are disordered and lack significant long-range contacts (Fig. S12 and S13A), and correspondingly motions on the intermediate timescale have the largest amplitudes (Figs. 5F and S10C). β-Synuclein is similar to α-synuclein, except that *A*_1_ is overestimated and becomes comparable to *A*_2_ (Fig. S8C). Aβ40 represents an intermediate scenario, where the N-half with extensive contacts has higher *A*_1_ than *A*_2_ but the C-half has comparable *A*_1_ and *A*_2_ (Fig. 5B).

Lastly let us take a closer look at the time constants and amplitudes of some residues noted above for elevated or depressed relaxation rates. The elevated *R*_2_ at Tyr39, Lys96, and Asp121 in α-synuclein (Fig. 3C) and at Lys311 and Val313 in tau K18 (Fig. S10A) can both be attributed to elevated *τ*_1_ (Figs. 5E and S10B), but the elevated *R*_2_ at Ala450 of SEV-NT (Fig. S9A) and at Tyr61-Phe64 and Asn79-Gln80 in A1-LCD (Fig. S11A) is due to elevated *A*_1_ (Figs. S9C and S11C). Many Gly residues have been seen to promote fast dynamics, as evidenced by dips in *R*_2_ and/or *R*_1_, including six in α-synuclein (Fig. 3C), five in β-synuclein (Fig. S8A), two in SEV-NT (Fig. S9A), four in tau K18 (Fig. S10A), and three in A1-LCD (Fig. S11A). All these Gly residues (or their immediate neighbors) can be recognized by dips in *A*_sum_ (Figs. 5F and S8C to S11C). In some cases the dips in *A*_sum_ are specifically due to dips in *A*_1_ (Figs. S9C to S11C) and occasionally are also be accompanied by dips in *τ*_1_ (Fig. S9B).

## Discussion

Based on experimentally validated MD simulations, we have characterized the sequence-dependent backbone dynamics of a variety of IDPs. The dynamics spans a wide range of timescales, from sub-ps to 10 ns, and the distribution of amplitudes on these timescales can vary greatly from IDP to IDP. Slow dynamics can be promoted by two mechanisms. The first mimics what happens in structured proteins, through the formation and packing of secondary structures, as exemplified by the helical C-half of the two HOX proteins. The second is through extensive local and long-range contacts, including hydrogen bonds, salt bridges, and cation- and amino-π interactions. Local contacts can rigidify a block of residues, as illustrated by the N-half of Aβ40. When long-range contacts are also formed, as typified by Al-LCD, the entire IDP becomes relatively compact and tumbles as a globule without any secondary structures. SEV-NT represents a hybrid of the above two mechanisms, where an α-helix is well-populated and also engages in long-range contacts.

Glycine is flexible and generally promotes fast dynamics. The flexibility allows glycine to play opposite roles in IDP dynamics. On the one hand, Gly repeats are very effective in interrupting rigid blocks, as illustrated by three GGGG motifs in A1-LCD and GGG and GG in SEV-NT. In tau K, four GGG motifs effectively break the IDP into four dynamically independent units. On the other hand, as seen in A1-LCD, Gly residues facilitate multiple modes of local interactions within rigid blocks and multiple modes of long-range contacts between rigid blocks.

We have fit NH time correlation functions to a sum of three exponentials. The three time constants fall in the ranges of 3-8 ns (*τ*_1_), 1-2 ns (*τ*_2_), and 0.1-0.2 ns (*τ*_3_) for all the IDPs. The greatest difference in dynamics between different regions of an IDP and among different IDPs is captured by the motional amplitudes of the different timescales. *A*_1_, the amplitude in the slow timescale, is dominant in the entire A1-LCD and SEV-NT, in the helical C-half of the two HOX proteins, and in the N-half of Aβ40, reflecting the effects of secondary structures and local and long-range interactions. In contrast, *A*_2_, the amplitude in the intermediate timescale, is either dominant or on par with *A*_1_ in the entire tau K18, α- and β-synuclein, in the disordered N-half of the two HOX proteins, and in the C-half of Aβ40, reflecting the lack of long-range or even local interactions. The *A*_1_- dominated IDPs or regions thereof also have a slightly higher *A*_sum_ than their *A*_2_- dominated counterparts, indicating that the suppression of slower dynamics in the former group extends all the way to the ultrafast (i.e., ps and sub-ps) timescale. Lastly the *A*_1_- dominated group has somewhat longer *τ*_1_ (and *τ*_2_ to a lesser extent) than the *A*_2_- dominated group.

It is instructive to compare the time constants and amplitudes of the IDPs with those of structured globular proteins. In the latter case the NH time correlation functions could be fit to the sum of two exponentials, with one time constant (*τ*_G_) for ns global tumbling and the other (*τ*_L_) for fast local motion ^29^. *τ*_G_ scales linearly with the number of residues (*τ*_G_ *≍* 0.051*N*) whereas *τ*_L_ fluctuates around 25 ps. The typical amplitudes of these two types of motions are 0.85 (*A*_G_) and 0.05 (*A*_L_). The *τ*_1_ motion of IDPs has some resemblance to the global tumbling of globular proteins, but differs in important ways. First, while the helical region of the two HOX proteins and the few most highly helical residues of SEV-NT have *A*_1_ amplitudes that approach the typical value of 0.85 for *A*_G_ (Figs. 5D, S7C, and S9C), other IDP residues, especially those in the IDPs that lack long-range contacts, have much lower *A*_1_ amplitudes. Second, the mean *τ*_1_ values of the IDPs lack dependence on sequence length. The expected *τ*_G_ value for a 40-residue protein like Aβ40 is 2.0 ns and that for a 140-residue like α-synuclein is 7.1 ns. However, Aβ40 has a mean *τ*_1_ of 3.8 ns and α-synuclein has a mean *τ*_1_ of 4.4 ns. Relative to the expected *τ*_G_, the higher mean *τ*_1_ of Aβ40 can be explained by its conformational ensemble being more open than the structure of a globular protein of the same sequence length, whereas the lower mean *τ*_1_ of α-synuclein can be rationalized if each of its residues belongs to a dynamically independent unit that is much shorter than the full-length protein. The fact that both Aβ40 and α-synuclein have a mean *τ*_1_ around 4 ns suggests that, for IDPs that lack well-populated secondary structures and long-range contacts, such dynamically independent units may be as short as 30 residues. As noted above, tau K18 may be divided into four such units demarcated by PGGG motifs, and the length of each unit is about 30 residues. So apparently two factors pull the *τ*_1_ values of IDPs in opposite directions: being more open than globular proteins could lead to higher *τ*_1_, but the accompanying loss of long-range contacts could lead to lower *τ*_1_.

The fast local motion seen in globular proteins with a very small amplitude does not show up in our three-exponential fits for IDPs, possibly being lumped into the missing amplitude. Instead, *τ*_2_ and *τ*_3_ emerge. An interesting question is motions on these timescales involve how many residues. If a 4-ns *τ*_1_ indeed involves 30 residues, then we could estimate a 1-ns *τ*_2_ could involve perhaps 5-7 residues, and a 0.15-ns *τ*_3_ might involve at most the nearest neighbors.

Sequence-dependent backbone dynamics can potentially code for functions or contribute to disease mechanisms, by endowing IDPs with versatile response to binding partners. In a previous study, based on sequence-dependent dynamics, we proposed that the rigid N-half of ChiZ-NT would be recalcitrant to interactions with binding partners whereas the flexible C-half would readily adapt to binding partners ^25^. This prediction was verified by a subsequent study on the association of ChiZ with acidic membranes, showing that the C-half is dominant in forming membrane contacts ^28^. A similar mechanism may be at work in the nucleation of Aβ40 fibrils, where the flexible C-half may readily undergo a conformation transition to form a β-sheet between monomers under a concentrated condition (Fig. 6A). Likewise tau K18 may utilize its flexibility to achieve multiple modes of binding with microtubule. On the other hand, the preformed three-helix bundle allows the HOX transcription factors to recognize their DNA target, while the dynamic N-terminal tail, which occasionally forms intramolecular contacts with the helix bundle in the free state, now binds to the co-transcription factor to provide further stabilization ^43^ (Fig. 6B). Similarly, the nascent α-helix in SEV-NT recognizes the target protein and becomes further rigidified on the target surface ^44^. Note that most of the local and long-range intramolecular contacts in free SEV-NT can still form in the bound state; a notable exception is the salt bridge between Glu429 and Arg482, as Arg482 now engages in intermolecular interactions with the target protein. It is also likely that many intramolecular contacts of A1-LCD become intermolecular under a concentrated condition, thereby enabling condensate formation ^17^. Intermolecular interactions in turn dictate dynamic properties of biomolecular condensates ^45^. Knowledge of sequence-dependent dynamics and its origins provides deep insight into functional and disease mechanisms.

**Figure 6.**
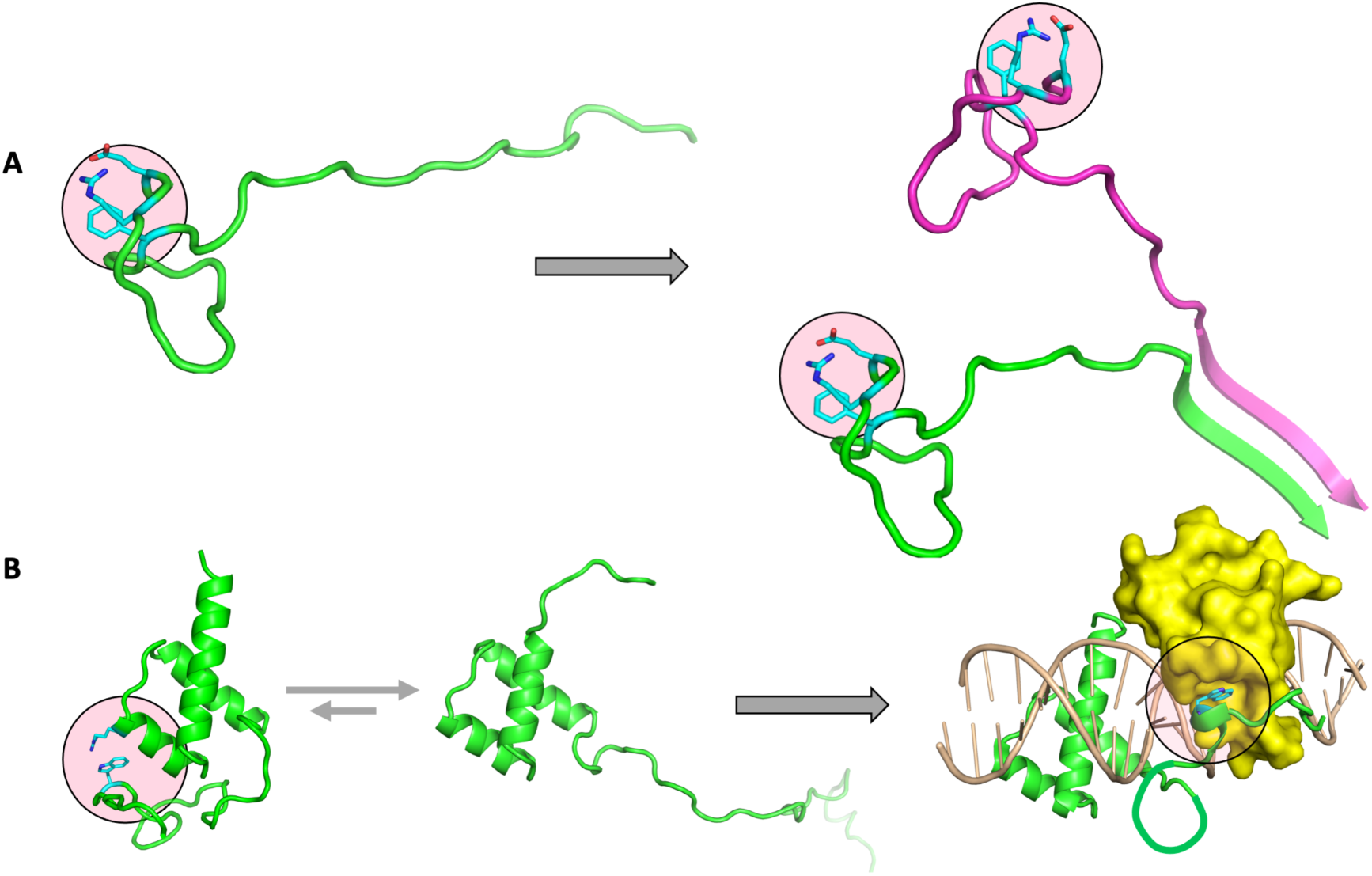
Mechanisms by which sequence-dependent backbone dynamics of IDPs codes for functions or contributes to diseases. (A) The fast dynamics of the C-half of Aβ40 allows it to readily form a β-sheet between monomers and nucleate fibrillation. (B) The rigid three-helix bundle of a HOX transcription factor allows it to recognize the DNA target; the dynamic N-terminal region can form intramolecular contacts in the free state but engage with the co-transcription factor to further stabilize the complex.

## Computational Methods

### Molecular Dynamics Simulations

MD simulations were run in AMBER18 ^4637^ using the ff14SB force field ^26^ for proteins and TIP4P-D for water ^27^. The eight IDPs span a wide range in sequence length and level of secondary structure. Accordingly we generated their initial structures in several ways. Those of Aβ40 were snapshots, with a short *α*-helix or a short *β*-sheet, from a previous simulation using a different force field^47^. Initial structures of HOX-SCR and HOX-DFD were generated by the I-TASSER web server ^48^; that of tau K18 was from the RaptorX web server ^49^. The initial structure of SEV-NT consisted of an *α*-helix for residues 476-492 [built in Pymol (https://pymol.org/)] and disordered N- and C-terminal regions (built by *tleap*). Initial structures of A1-LCD and *α*- and *β*-synuclein were generated by the TraDES web server ^50^. Each initial structure was placed into a rectangular box, with a solvent layer ranging from ∼20 Å (for the more open) to ∼50 Å (for the more compact). Na^+^ and Cl^-^ were added to neutralize the systems and provide the experimental salt concentrations. The initial extent of secondary structures, total number of atoms, NaCl concentration, and simulation temperature for each IDP are listed in Table S1.

For each system, with *sander*, energy minimization (2000 steps of steepest descent and 3000 steps of conjugate gradient) was followed by heating from 0 to the final temperature (Table S1) over 100 ps at a 1 fs timestep. Temperature was regulated by the Langevin thermostat ^41^ with a 3.0 ps^-1^ damping constant. Bond lengths involving hydrogens were constrained using the SHAKE algorithm ^51^. Long-range electrostatic interactions were treated by the particle mesh Ewald method ^52^. The cutoff distance for nonbonded interactions were 10 Å. The simulations then continued on GPUs using *pmemd.cuda* ^53^ in four replicates with different random seeds at constant temperature and pressure (1 atm) at a 2 fs timestep, initially for 3 ns and then for 3.2 μs. In the case of Aβ40 and A1-LCD, four replicates were prepared from the beginning using different initial structures. Pressure was regulated using the Berendsen barostat ^42^. The final 2.5 μs of each trajectory, with snapshots saved every 20 ps, was used for analysis.

### Chemical shifts, Small Angle X-ray Scattering Profiles, and Paramagnetic Relaxation Enhancements

These properties were averaged over MD snapshots saved every 1 ns. Chemical shifts were calculated using SHIFTX2 ^54^. Secondary Cα and Cβ chemical shifts were calculated by subtracting random-coil values from POTENCI ^55^. SAXS profiles were calculated using FoXS ^56^, with calculated profile for each snapshot scaled to optimize the match with the experimental data. PREs were calculated using DEER-PREdict ^57^.

### Secondary Structures

Secondary structures were calculated using the *dssp* command in *cpptraj* ^58^. These results were expanded to include PPII formation when three or more consecutive coil residues fell into the PPII region of the Ramachandran map.

### Contact Numbers and Contact Maps

Distances between heavy atoms (except those in the same residue) were calculated by loading trajectories into MDTraj ^59^. A contact was defined when two heavy atoms were within 3.5 Å of each other. The contact number of a residue was the number of residues beyond the immediate neighbors with which at least one contact was formed. This number was calculated for each snapshot and then averaged over all the saved snapshots. For calculating the contact map, the contact frequency between any two heavy atoms from two different side chains was calculated by counting the fraction of snapshots in which that contact was formed.

### NMR Relaxation Properties

For each non-proline residue, the time correlation function of the NH bond vector was calculated as

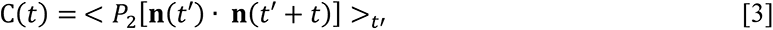

where **n**(*t*′) is the unit vector along the NH bond at time *t*′; *P*_2_(*x*) is the second order Legendre polynomial; and < ⋯ >_t′_ denotes time average over each trajectory. After further averaging over the four replicate simulations, the correlation function was fit to a tri-exponential function,

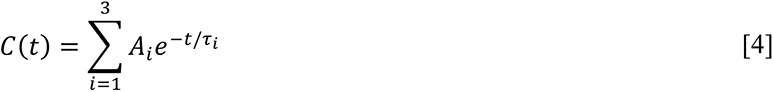

by a custom python code ^25^. The fit was done without constraints on any parameters. In particular, the sum of the amplitudes was not constrained to 1, allowing for missing amplitudes of ultrafast motions (i.e., faster than 20 ps at which snapshots were saved). The default for the upper bound of the time range was 15 ns, but for a small number of residues fitting errors were reduced by increasing this upper bound. Following our previous study ^29^, the three time constants were corrected by a scaling factor according to Eq [2] in order to remove dynamic distortions of the Langevin thermostat used in the MD simulations.

The spectral density for the time correlation function of Eq [4] was

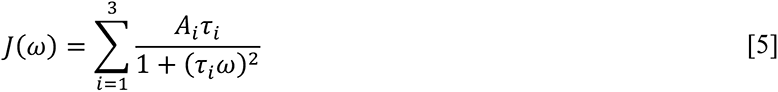

Finally *R*_1_, *R*_2_, and NOE were given by

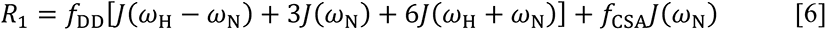

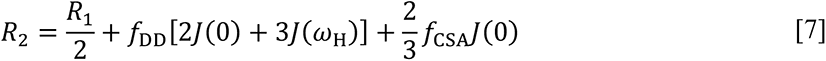

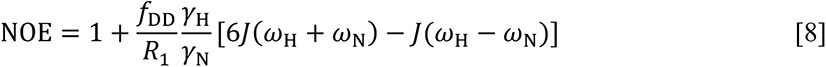

where 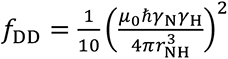 and 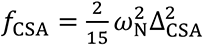. The meanings of the symbols are: *μ*_0_, permittivity of free space; *ħ*, reduced Plank constant; γ_H_ and γ_N_, gyromagnetic ratios of hydrogen and nitrogen; *ω*_H_ = γ_H_*B*_0_, Larmor frequency of hydrogen; *ω*_N_, counterpart of nitrogen; *r*_NH_, NH bond length (set at 1.02 Å); and Δ_678_ (= –170 ppm), chemical shift anisotropy of nitrogen. The magnetic field strengths were 600 MHz for Aβ40 and α-and β-synuclein, 700 MHz for tau K18, 800 MHz for HOX-DFD, HOX-SCR, and A1-LCD, 850 MHz for SEV-NT. RMSEs for *R*_1_, *R*_2_, and NOE were calculated over non-terminal residues.

## Acknowledgments

This work was supported by National Institutes of Health Grant R35 GM118091. We thank Alan Hicks and Ramash Prasad for technical assistance.

## Author contributions

S.D, M.M., and H.-X.Z. designed and performed the research, analyzed the data, and wrote the manuscript.

## Competing financial interests

The authors declare no competing financial interests.

## Notes

### Competing Interest Statement

The authors have declared no competing interest.

