## Supporting Tables and Figures for "Sequence-Dependent Backbone Dynamics of Intrinsically Disordered Proteins"

Supporting Information

Table S1. Simulation and validation details of eight IDPs.

| IDP name | starting structure | # of atoms | NaCl (mM) | simulation temp (K) | exp data for validation[a] |
|---|---|---|---|---|---|
| Aβ40 | short α-helix or β-sheet | 240733 - 245263 | 50 | 278 | CS [1], inter-residue NOE [1], relaxation [2] |
| HOX-SCR | 3 α-helices | 276948 | 200 | 300 | CS [3], relaxation [3] |
| HOX-DFD | 3 α-helices | 288806 | 200 | 300 | CS [3], relaxation [3] |
| SEV-NT | 1 α-helix | 356926 | 500 | 274 | CS [4], relaxation [4] |
| tau K18 | disordered | 276036 | 100 | 300 | SAXS [5], CS [6], relaxation [6] |
| A1-LCD | disordered | 310922 - 313872 | 50 | 288 | SAXS [7], CS [7], relaxation [7] |
| β-synuclein | disordered | 360102 | 100 | 300 | CS [8], PRE [9], relaxation [8] |
| α-synuclein | disordered | 371305 | 100 | 300 | SAXS [10], CS [11], PRE [9, 12-14], relaxation [15] |

[a]Abbreviations: CS, chemical shift; NOE, nuclear Overhauser effect; PRE, paramagnetic relaxation enhancement; SAXS, small-angle X-ray scattering

Table S2. Mean values and standard deviations of time constants and amplitudes of eight IDPs.

| IDP name | $\tau_1$ (ns) | $\tau_2$ (ns) | $\tau_3$ (ns) | $A_1$ | $A_2$ | $A_3$ | $A_{sum}$ |
|---|---|---|---|---|---|---|---|
| Aβ40 | 3.79 ± 0.68 | 1.12 ± 0.31 | 0.17 ± 0.05 | 0.36 ± 0.12 | 0.33 ± 0.07 | 0.15 ± 0.05 | 0.85 ± 0.02 |
| HOX-SCR | 5.81 ± 0.84 | 1.59 ± 0.56 | 0.17 ± 0.06 | 0.59 ± 0.23 | 0.18 ± 0.11 | 0.09 ± 0.09 | 0.87 ± 0.04 |
| HOX-DFD | 5.76 ± 1.01 | 1.53 ± 0.41 | 0.15 ± 0.05 | 0.59 ± 0.24 | 0.18 ± 0.11 | 0.09 ± 0.09 | 0.86 ± 0.05 |
| SEV-NT | 6.86 ± 1.02 | 1.58 ± 0.40 | 0.21 ± 0.06 | 0.57 ± 0.15 | 0.20 ± 0.08 | 0.10 ± 0.05 | 0.87 ± 0.03 |
| tau K18 | 3.94 ± 0.71 | 1.03 ± 0.27 | 0.17 ± 0.05 | 0.23 ± 0.06 | 0.36 ± 0.04 | 0.22 ± 0.05 | 0.81 ± 0.04 |
| A1-LCD | 5.93 ± 0.50 | 1.30 ± 0.30 | 0.16 ± 0.04 | 0.55 ± 0.11 | 0.18 ± 0.05 | 0.11 ± 0.06 | 0.84 ± 0.03 |
| β-synuclein | 5.22 ± 0.70 | 1.27 ± 0.28 | 0.19 ± 0.04 | 0.32 ± 0.08 | 0.32 ± 0.04 | 0.19 ± 0.04 | 0.82 ± 0.02 |
| α-synuclein | 4.43 ± 0.72 | 1.05 ± 0.19 | 0.17 ± 0.04 | 0.24 ± 0.06 | 0.35 ± 0.04 | 0.22 ± 0.04 | 0.81 ± 0.03 |

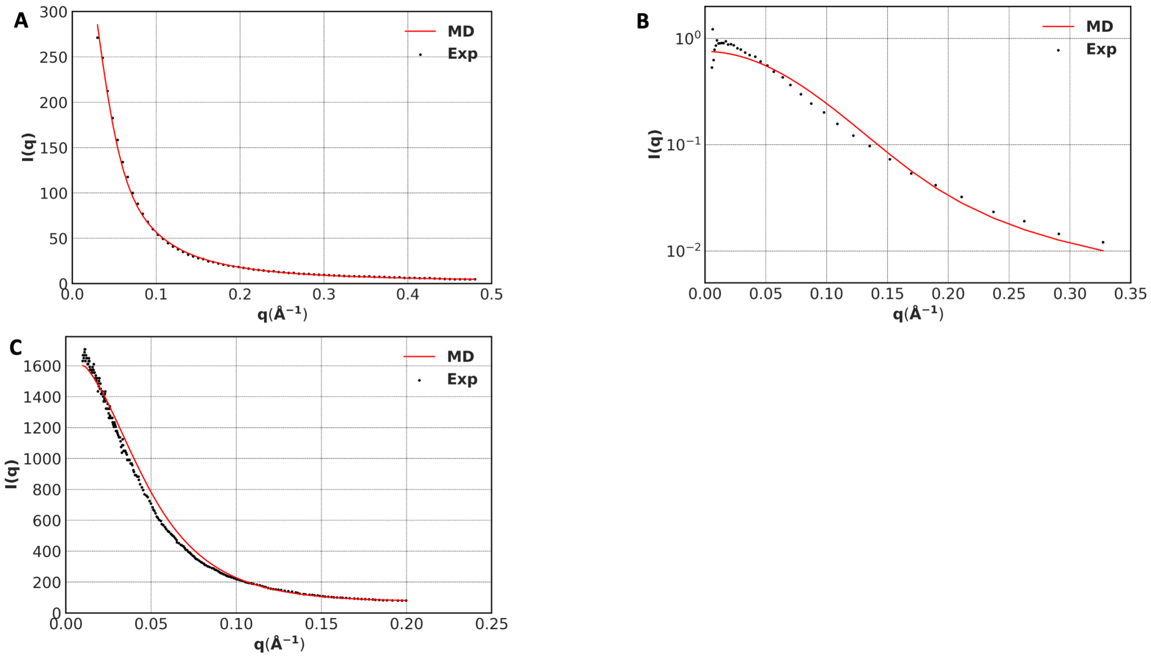


**Figure S1.** Comparison of SAXS profiles calculated from MD simulations and their experimental counterparts. (A) tau K18. (B) A1-LCD. (C) α-synuclein.

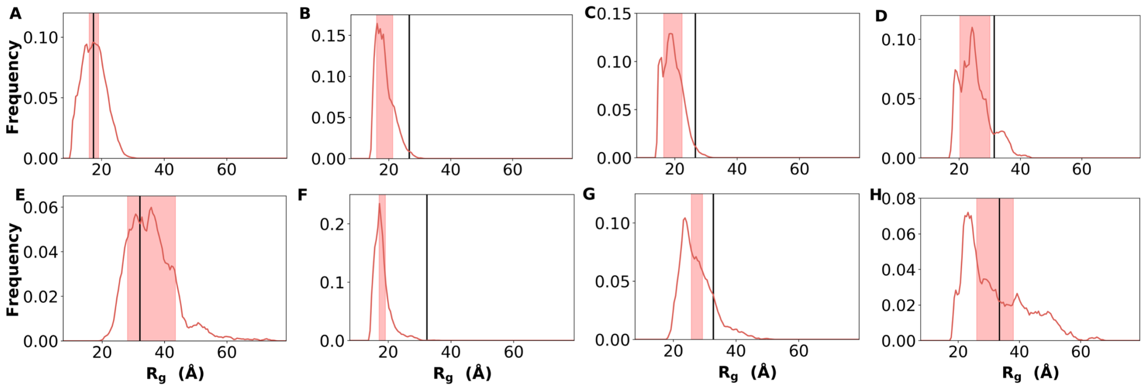


**Figure S2.** Histograms of the radii of gyration ($R_g$) of the eight IDPs. (A) Aβ40. (B) HOX-SCR. (C) HOX-DFD. (D) SEV-NT. (E) tau K18. (F) A1-LCD. (G) β-synuclein. (H) α-synuclein. Red bands display mean ± standard deviation; black vertical lines represent $R_g$ values predicted according to the scaling relation in Eq [1].

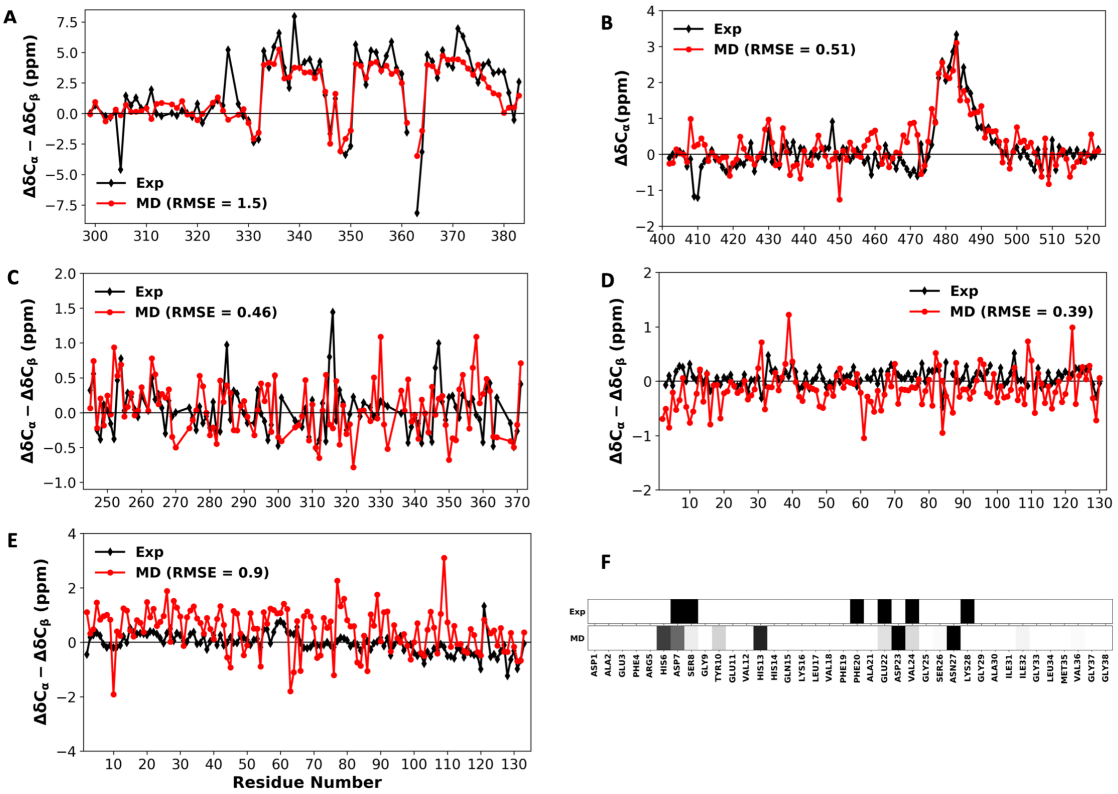


**Figure S3.** Experimental validation of MD conformational ensembles. (A-E) Comparison of calculated and experimental secondary chemical shifts for HOX-SCR, SEV-NT, tau K18, A1-LCD, and β-synuclein. RMSE values are shown in the legends. (F) Comparison of experimental (top, black bars) and calculated (bottom) αN($i$, $i$ + 2) NOEs for Aβ40. The calculated results display the effective interproton distance, $< r^{-6} >^{-1/6}$, where $r$ is the distance between $H_\alpha$ at residue $i$ and $H_N$ at residue $i$ + 2, as a bars on a gray scale, with values at 4.2 Å or blow in black and values at 4.5 Å or above in white.

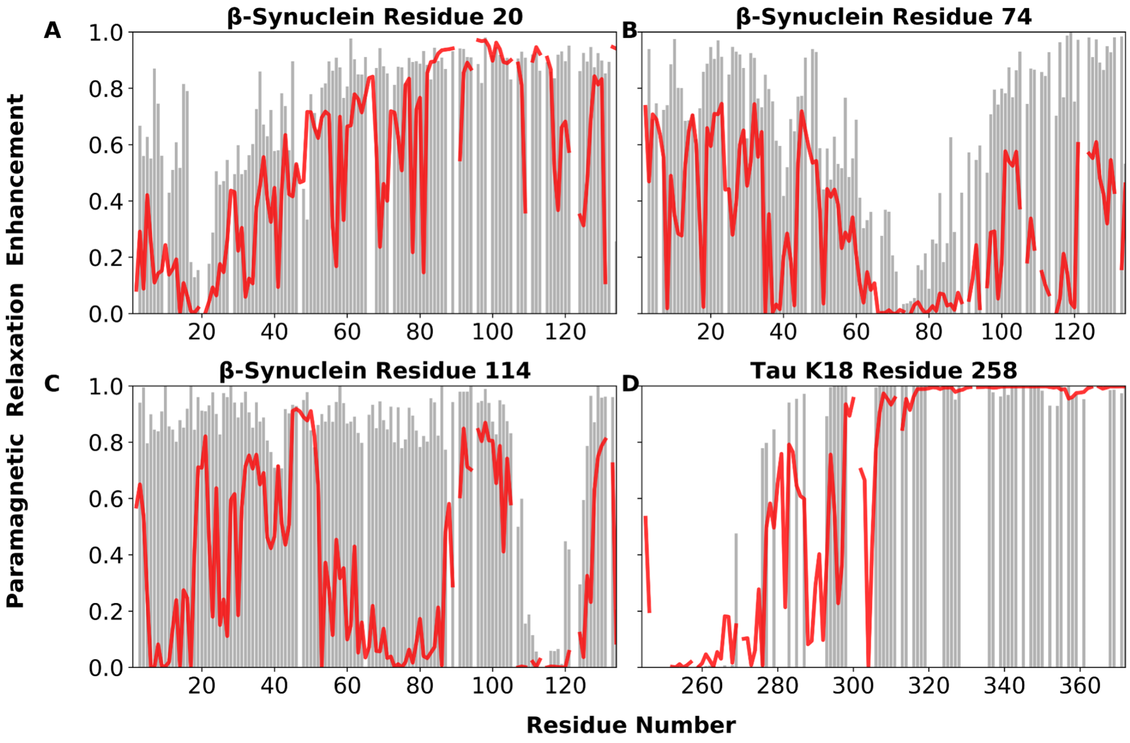


**Figure S4.** Comparison of calculated (red curves) and experimental (gray bars) PREs. (A-C) Results for spin labels at three residues in β-synuclein. (D) Results for a spin label in tau K18.

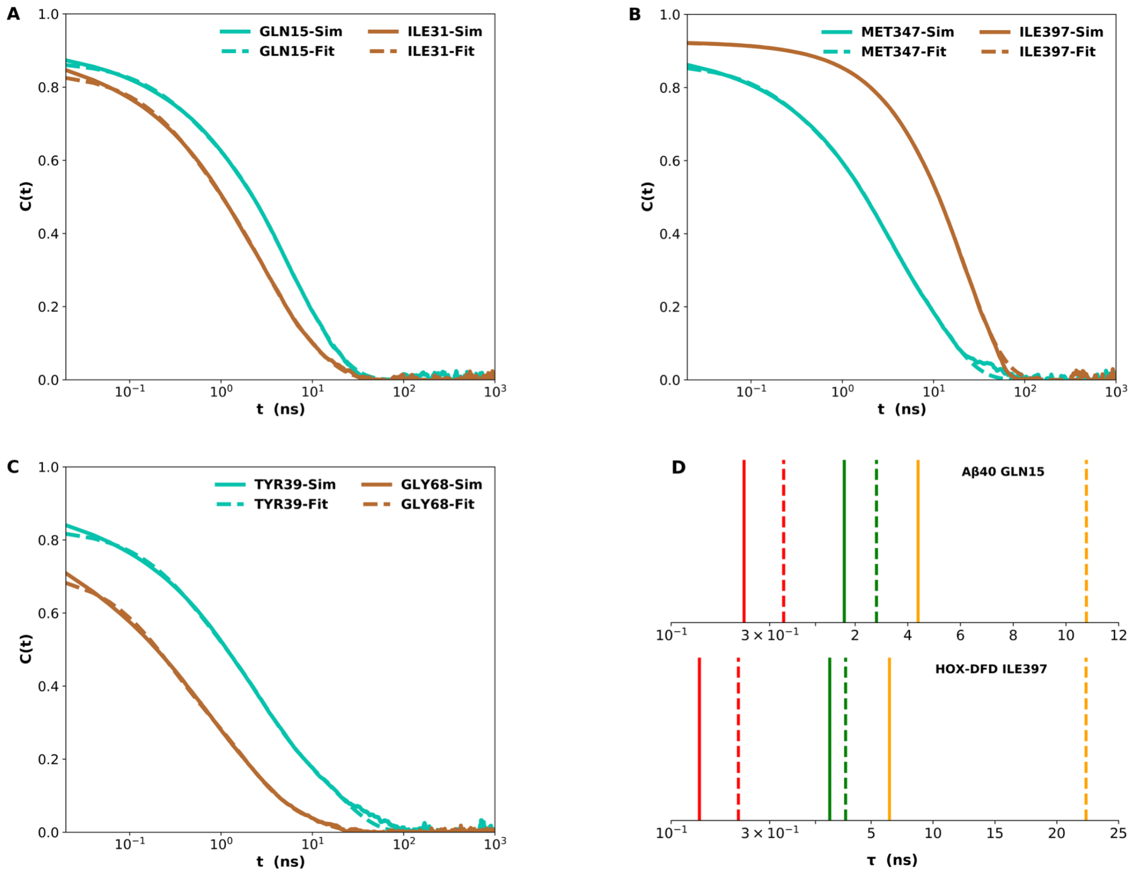


**Figure S5.** Illustrative fits of NH time correlation functions to a sum of three exponentials and correction of the time constants to remove dynamic distortion by the Langevin thermostat. (A-C) Fits of two residues each from Aβ40, HOX-DFD, and α-synuclein. (D) Raw time constants (dashed vertical lines) and corrected values according to Eq [2] for two residues. $\tau_1$, $\tau_2$, and $\tau_3$ are displayed in orange, green, and red, respectively.

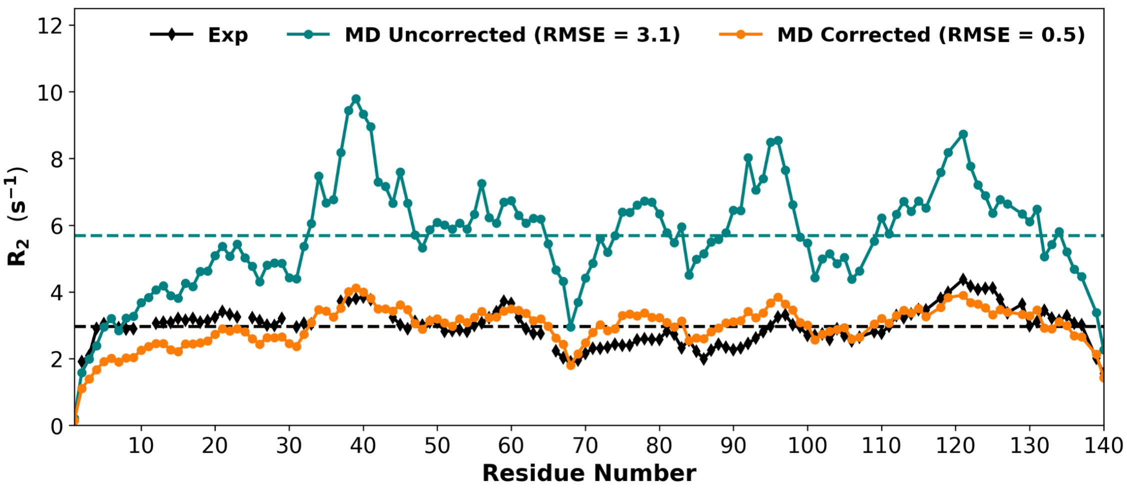


**Figure S6.** Transverse relaxation rates calculated with uncorrected time constants and with time constants after correction according to Eq [2], for α-synuclein. The experimental results are also shown (black). RMSEs of uncorrected and corrected calculations are given in the legend.

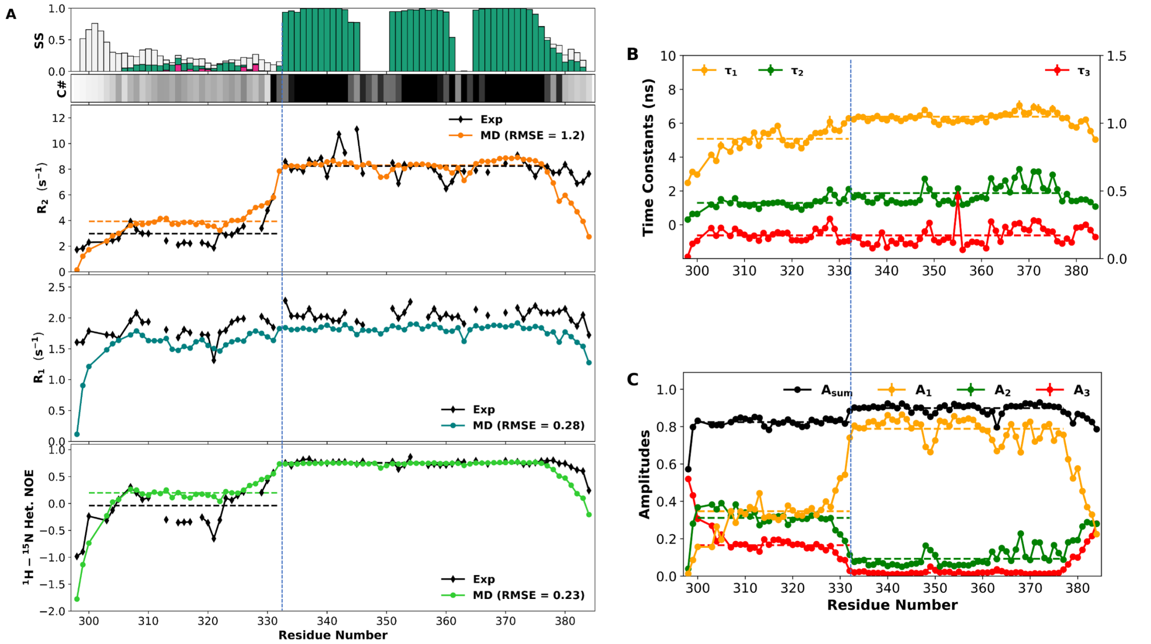


**Figure S7.** Residue-specific backbone dynamics in HOX-SCR. (A) Secondary structure propensities, average contact numbers, and calculated and experimental $R_2$, $R_1$, and NOE. Horizontal dashed lines show average values of $R_2$ and NOE over the N-half and C-half (demarcated by black vertical lines). (B) The three time constants of the fits of the NH time correlation functions to a sum of three exponentials. The left ordinate displays the scale for $\tau_1$ and $\tau_2$ whereas the right ordinate displays the scale for $\tau_3$. Horizontal dashed lines show average values over the N-half and C-half (demarcated by a black vertical line, for $\tau_1$ and $\tau_2$) or over the entire sequence (for $\tau_3$). (C) The amplitudes of the three exponentials and their sum. Horizontal dashed lines show average values over the N-half and C-half (demarcated by black vertical line).

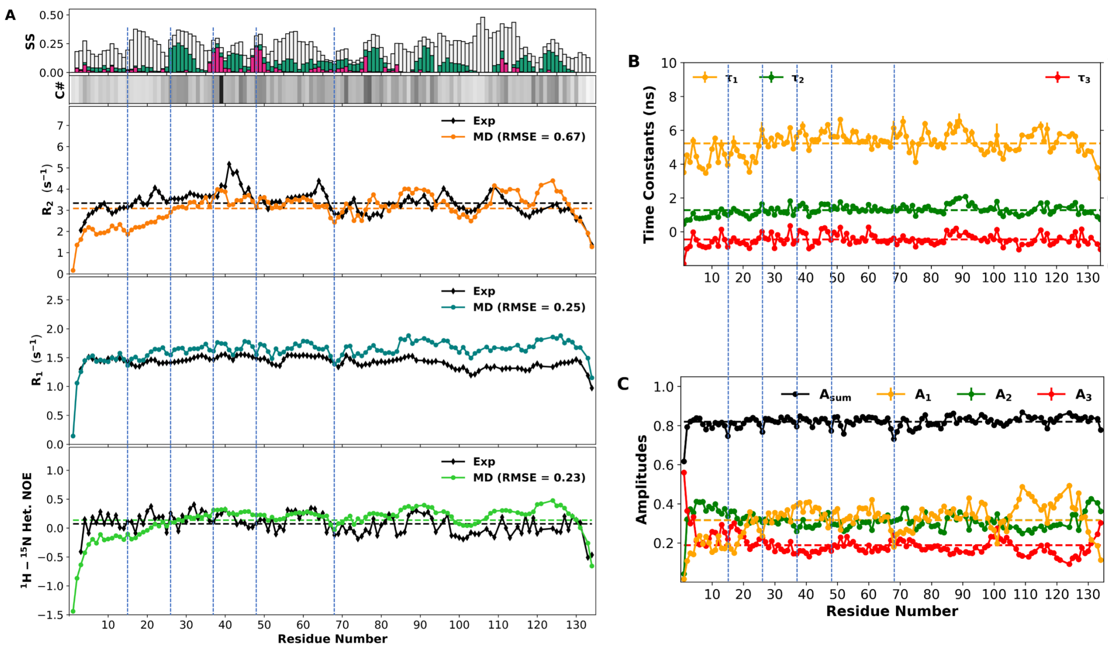


**Figure S8.** Residue-specific backbone dynamics in β-synuclein. (A) Secondary structure propensities, average contact numbers, and calculated and experimental $R_2$, $R_1$, and NOE. The calculated $R_2$ was scaled down by a factor of 1.25 to account for a 12 K higher temperature in the MD simulations than in the NMR relaxation experiment [8]. Horizontal dashed lines show average values of $R_2$ and NOE over the entire sequence. Blue vertical lines are placed at residues 15, 26, 37, 48, and 68 to indicate dips in $R_1$ (and $R_2$ in the case of Gly68). (B) The three time constants of the fits of the NH time correlation functions to a sum of three exponentials. The left ordinate displays the scale for $\tau_1$ and $\tau_2$ whereas the right ordinate displays the scale for $\tau_3$. Horizontal dashed lines show average values over the entire sequence. (C) The amplitudes of the three exponentials and their sum. Horizontal dashed lines show average values over the entire sequence. Blue vertical lines are placed at residues 15, 26, 37, 48, and 68 to indicate dips in $A_{\text{sum}}$.

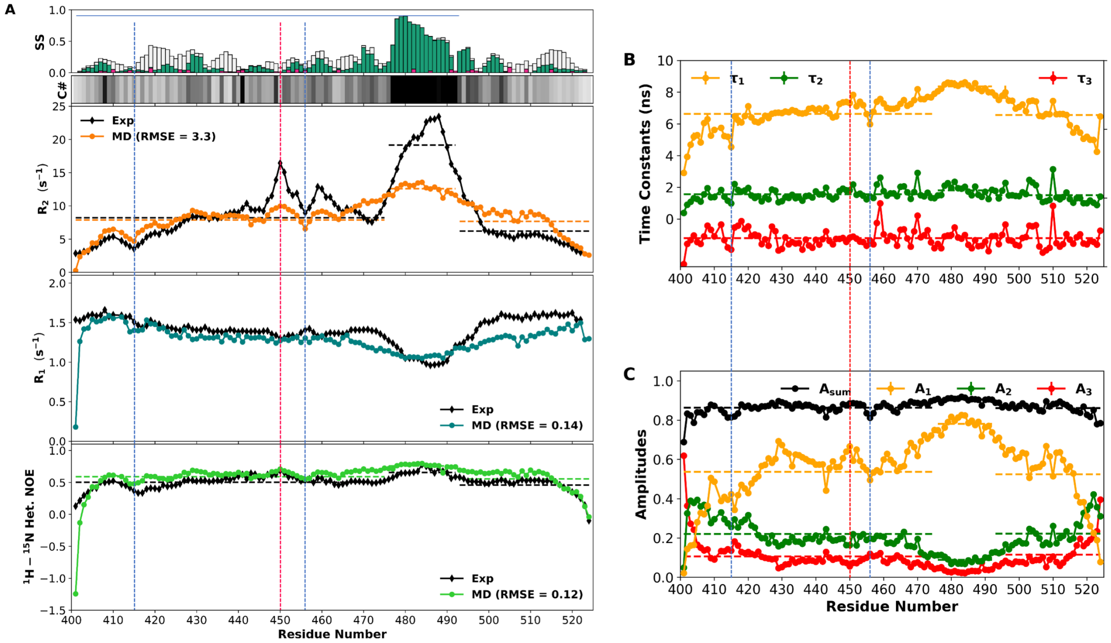


**Figure S9.** Residue-specific backbone dynamics in SEV-NT. (A) Secondary structure propensities, average contact numbers, and calculated and experimental $R_2$, $R_1$, and NOE. Horizontal dashed lines show average values of $R_2$ and NOE over the N-terminal, helical (residues 476-492), and C-terminal regions. Blue vertical lines are placed at Gly415 and Gly456 to indicate dips in $R_2$ and NOE; a red vertical line is placed at Ala450 to indicated elevated $R_2$. (B) The three time constants of the fits of the NH time correlation functions to a sum of three exponentials. The left ordinate displays the scale for $\tau_1$ and $\tau_2$ whereas the right ordinate displays the scale for $\tau_3$. Horizontal dashed lines show average values over the N-terminal, helical, and C-terminal regions (for $\tau_1$ and $\tau_2$) or over the entire sequence (for $\tau_3$). (C) The amplitudes of the three exponentials and their sum. Horizontal dashed lines show average values over the N-terminal, helical, and C-terminal regions. Blue vertical lines are placed at Gly415 and Gly456 to indicate dips in $\tau_1$, $A_1$, and $A_{sum}$; a red vertical line is placed at Ala450 to indicated elevated $A_1$.

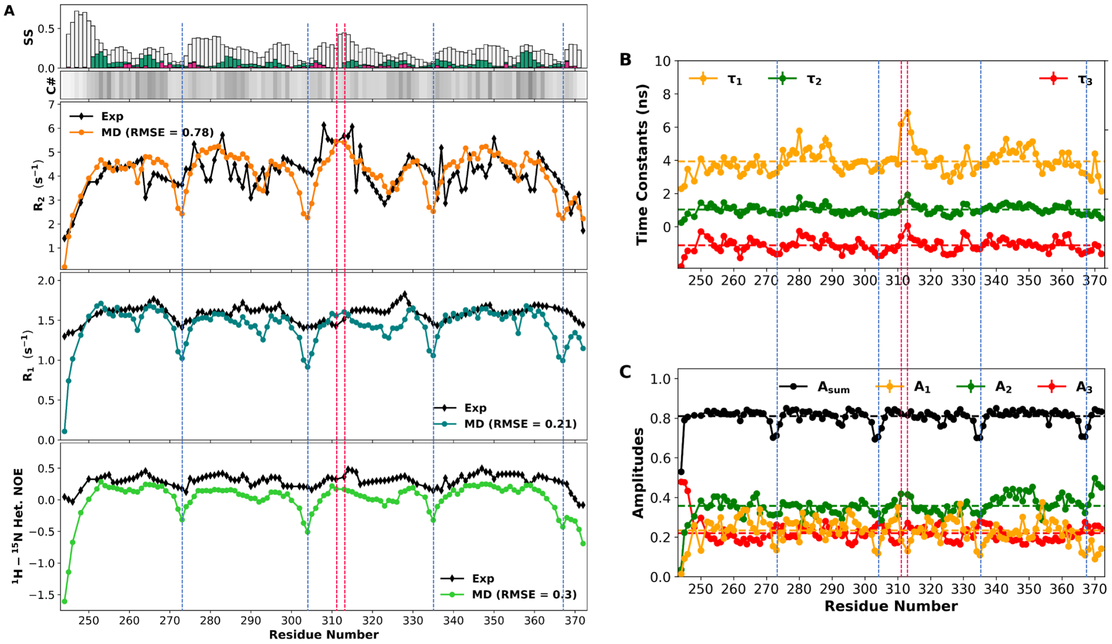


**Figure S10.** Residue-specific backbone dynamics in tau K18. (A) Secondary structure propensities, average contact numbers, and calculated and experimental $R_2$, $R_1$, and NOE. The calculated $R_2$ was scaled down by a factor of 1.5 to account for a 17 K higher temperature in the MD simulations than in the NMR relaxation experiment [6]. Blue vertical lines are placed at Gly273, 304, 335, and 367 to indicate significant dips in $R_2$, $R_1$, and NOE; red vertical lines are placed at Lys311 and Val313 to indicated elevated $R_2$. (B) The three time constants of the fits of the NH time correlation functions to a sum of three exponentials. The left ordinate displays the scale for $\tau_1$ and $\tau_2$ whereas the right ordinate displays the scale for $\tau_3$. Horizontal dashed lines show average values over the entire sequence. Red vertical lines are placed at Lys311 and Val313 to indicated elevated $\tau_1$. (C) The amplitudes of the three exponentials and their sum. Horizontal dashed lines show average values over the entire sequence. Blue vertical lines are placed at Gly273, 304, 335, and 367 to indicate dips in $A_1$ and $A_{sum}$.

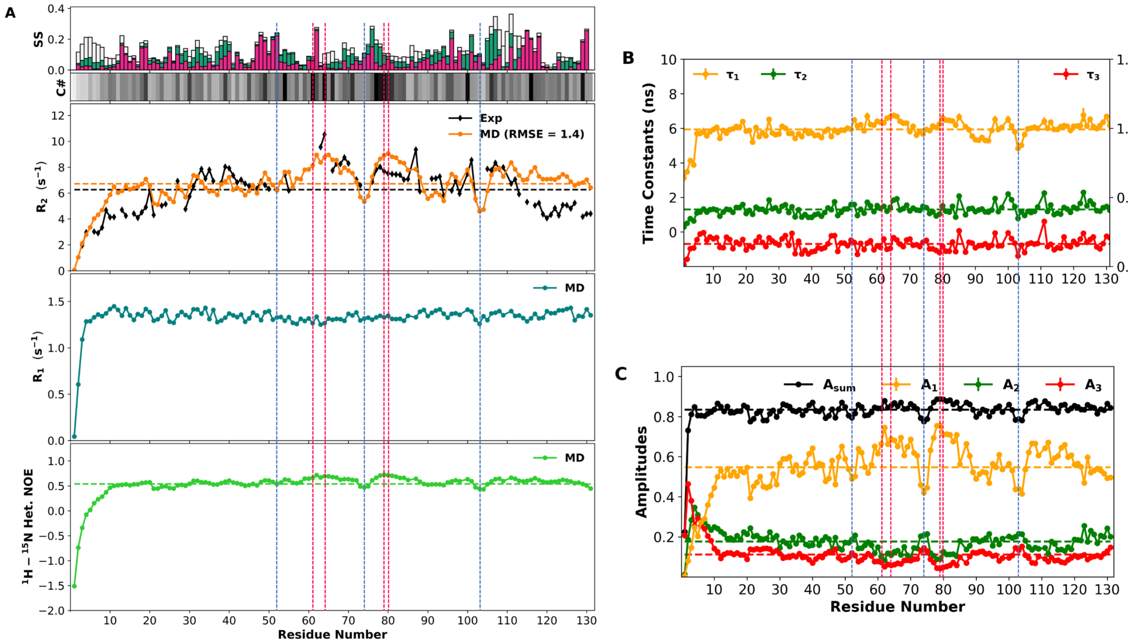


**Figure S11.** Residue-specific backbone dynamics in A1-LCD. (A) Secondary structure propensities, average contact numbers, calculated $R_2$, $R_1$, and NOE, and experimental $R_2$. Horizontal dashed lines show average values of $R_2$ and NOE over the entire sequence. Blue vertical lines are placed at Gly52, 74, and 103 to indicate dips in $R_2$ and NOE; red vertical lines are placed at Tyr61, Phe64, Asn79, and Gln80 to indicated elevated $R_2$. (B) The three time constants of the fits of the NH time correlation functions to a sum of three exponentials. The left ordinate displays the scale for $\tau_1$ and $\tau_2$ whereas the right ordinate displays the scale for $\tau_3$. Horizontal dashed lines show average values over the entire sequence. (C) The amplitudes of the three exponentials and their sum. Horizontal dashed lines show average values over the entire sequence. Blue vertical lines are placed at Gly52, 74, and 103 to indicate dips in $A_1$ and $A_{\text{sum}}$; red vertical lines are placed at Tyr61, Phe64, Asn79, and Gln80 to indicated elevated $A_1$.

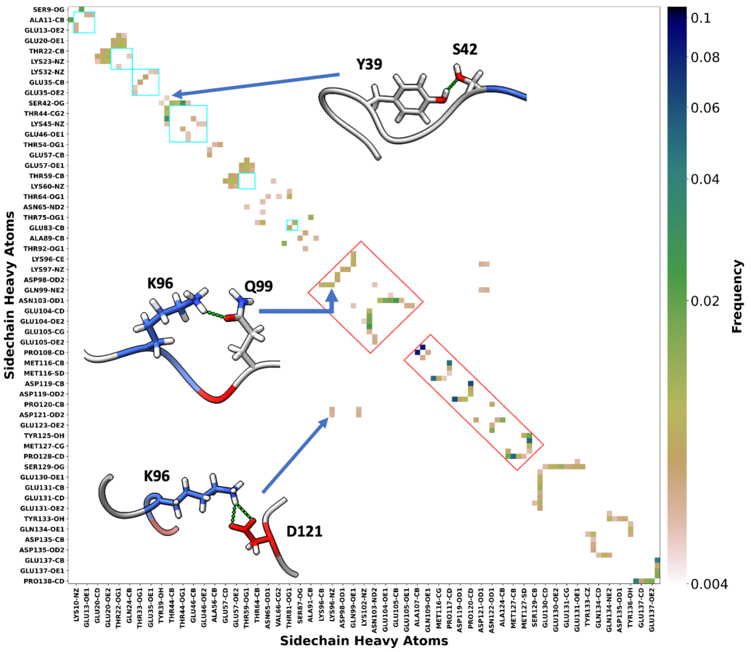


**Figure S12.** Side chain-side chain contact maps for α-synuclein. Contacts with frequencies higher than 0.005 are shown. Blocks of residues with tendencies for extensive contacts are highlighted by red boxes. KTKE motifs are identified by cyan boxes. Insets: snapshots illustrating selected contacts.

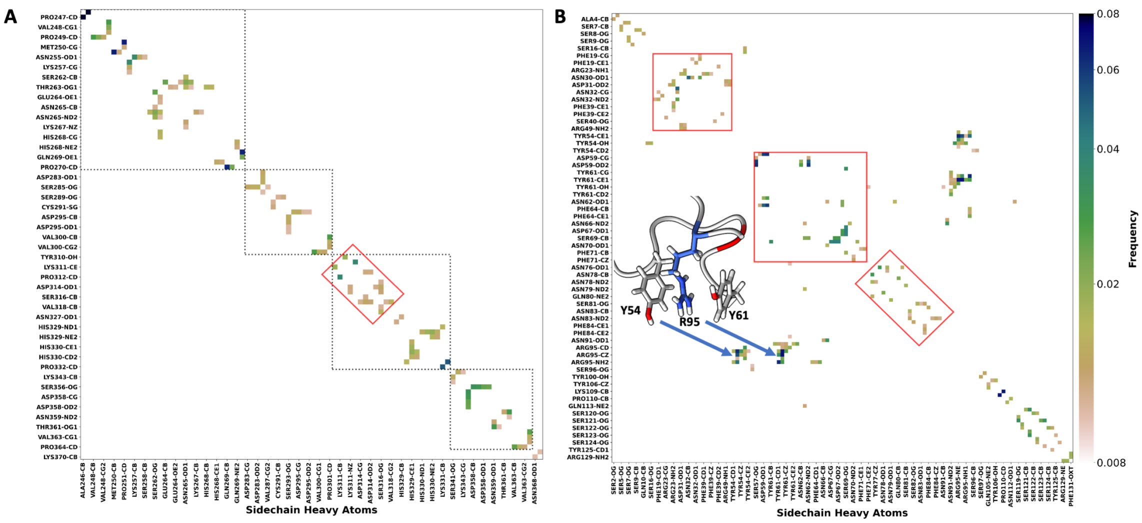


**Figure S13.** Side chain-side chain contact maps. (A) tau K18. Black dashed boxes indicate four repeats ending with PGGG motifs. (B) A1-LCD. Contacts with frequencies higher than 0.01 are shown. Blocks of residues with tendencies for extensive contacts are highlighted by red boxes. Inset: snapshot illustrating selected contacts.

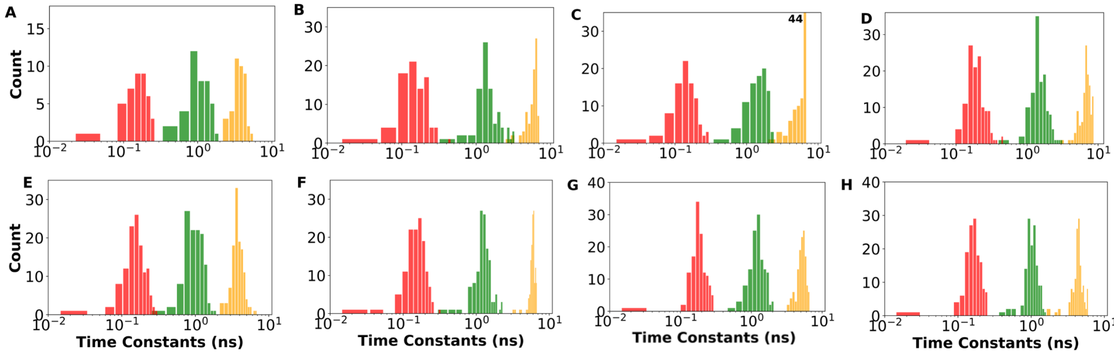


**Figure S14.** Histograms of the three time constants (on a log scale) for each of the eight IDPs. (A) Aβ40. (B) HOX-SCR. (C) HOX-DFD. (D) SEV-NT. (E) tau K18. (F) A1-LCD. (G) β-synuclein. (H) α-synuclein. Results for $\tau_1$, $\tau_2$, $\tau_3$ are shown in orange, green, and red, respectively.
